# Mef2c Controls Postnatal Callosal Axon Targeting by Regulating Sensitivity to Ephrin Repulsion

**DOI:** 10.1101/2025.01.22.634300

**Authors:** Sriram Sudarsanam, Luis Guzman-Clavel, Nyle Dar, Jakub Ziak, Naseer Shahid, Xinyu O. Jin, Alex L. Kolodkin

## Abstract

Cortical connectivity is contingent on ordered emergence of neuron subtypes followed by the formation of subtype-specific axon projections. Intracortical circuits, including long-range callosal projections, are crucial for information processing, but mechanisms of intracortical axon targeting are still unclear. We find that the transcription factor Myocyte enhancer factor 2-c (Mef2c) directs the development of somatosensory cortical (S1) layer 4 and 5 pyramidal neurons during embryogenesis. During early postnatal development, *Mef2c* expression shifts to layer 2/3 callosal projection neurons (L2/3 CPNs), and we find a novel function for *Mef2c* in targeting homotopic contralateral cortical regions by S1-L2/3 CPNs. We demonstrate, using functional manipulation of EphA-EphrinA signaling in *Mef2c-*mutant CPNs, that Mef2c downregulates *EphA*6 to desensitize S1-L2/3 CPN axons to EphrinA5-repulsion at their contralateral targets. Our work uncovers dual roles for *Mef2c* in cortical development: regulation of laminar subtype specification during embryogenesis, and axon targeting in postnatal callosal neurons.

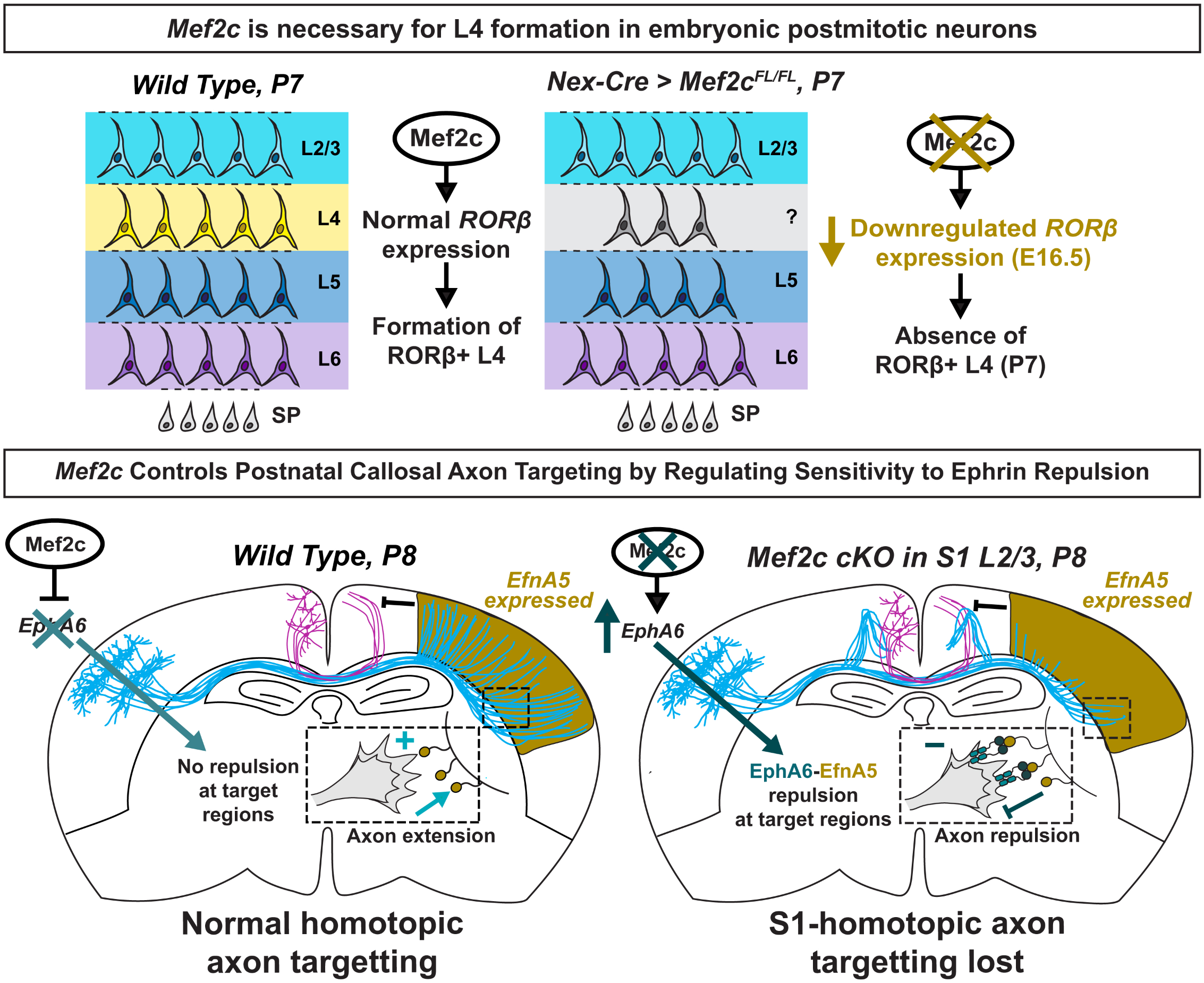

**HIGHLIGHTS:**

- Mef2c is required for the development of L4 and L5 neurons in the embryonic neocortex
- Postnatally, *Mef2c* is enriched in L2/3 neurons and is required for axon targeting
- L2/3-specific *Mef2c* deletion leads to *EphA6* upregulation
- *Mef2c* deletion in L2/3 neurons sensitizes them to EfnA5 repulsion in the contralateral cortex

## INTRODUCTION

Ordered connectivity in the brain enables transformation of sensory stimuli into meaningful perceptions, which in turn guides learning, memory and decision making. In the mammalian brain, sensory integration and transformation primarily occur in the cerebral cortex, mediated in part by long-range connections between cortical areas. Layer 2/3 of the murine primary somatosensory cortex (S1) is home to a major population of long-range intracortical projection neurons^1,2^. These neurons elaborate contralateral projections through the corpus callosum, the largest white-matter tract in the mammalian brain, to mediate interhemispheric communication^3^. Corpus callosum development proceeds in a stepwise fashion^3–6^, and recent advances have identified callosal neuron-specific transcriptional regulators^7,8,9,10^ as well as extrinsic signals^11,12^, which together regulate the initial channeling of axons into the white matter and promote subsequent CNS midline crossing. Midline crossing is followed by topographically organized target innervation, whereby callosal neurons innervate contralateral cortical domains that match their region of origin in the ipsilateral cortex. Precise homotopic targeting of callosal projections is crucial for information flow between cortical areas, and deficits in callosal connectivity are associated with neurodevelopmental^13,14,15^ and neuropsychiatric^16^ disorders. While there is some evidence for axon-axon interactions directing homotopic callosal axon targeting^17–19^, the molecular determinants that operate in callosal neurons and in their targets to promote appropriate axon targeting remain elusive.

The Myocyte enhancer factor 2 (Mef2) family of four paralogous transcription factors controls development and differentiation across different tissues^20^. Of these, Mef2c regulates several aspects of neuronal development, including neurogenesis and differentiation^21–25^, neuronal survivial^26–28^, axon and dendrite elaboration^23,24,29,30^ and synaptic development^31–37^. Loss-of-function (LOF) mutations and deficiencies in *MEF2C* are associated with conditions such as autism spectrum disorder, intellectual disability, epilepsy and schizophrenia^38,39^. *Mef2c* is broadly expressed in the cerebral cortex during embryogenesis^40–42^ and regulates the development of cortical laminar organization^21^. Early broad expression of *Mef2c* in the embryonic cortex transitions during postnatal development to laminar enrichment in superficial Layer 2/3 callosal projection neurons (L2/3 CPNs)^41^, raising the possibility of important Mef2c functions in callosal axon projection and targeting. Furthermore, the expression of several axon-guidance genes is dysregulated following *Mef2c* deletion in the cortex^34,37^, but to date these alterations have not been functionally linked to specific defects in cortical connectivity.

Among the dysregulated genes in the *Mef2c* mutant cortex are several Class-A Ephrin receptors^34,37^. Ephrins and their receptors were initially characterized as repulsive regulators of axon guidance^43,44^ that underlie the formation of topographic maps^45^, and Ephrin-Eph signaling has also been implicated in many aspects of cortical development ^11,46–53^. Further, differential expression of the ligand *Ephrin A5* (*EfnA5)* in the developing cortex regulates area-specific innervation by axons from distinct thalamic nuclei, which differ in their levels of *EphA* receptor expression ^54–58^. Interestingly, L2/3 CPNs in S1 express low levels of *EphA* receptors^59,60^ and project to homotopic contralateral cortical domains that exhibit high *EfnA5* expression. This raises the question of whether differential expression of *EfnA5* in cortical areas is also functionally relevant for targeting intracortical axons.

Here, we first investigate the function of embryonic *Mef2c* in cortical organization following early pan-cortical deletion. We then explore postnatal roles for *Mef2c* in callosal neuron axon outgrowth and targeting. To bypass non-autonomous effects of disrupted cortical organization due to embryonic deletion of *Mef2c,* we employ *in utero* electroporation-based strategies for robust post-mitotic L2/3 CPN-specific sparse labeling and genetic manipulation. We assess candidate cell-surface molecules likely to be regulated by Mef2c that influence callosal neuron axon targeting and identify a role for EphrinA5-EphA signaling, downstream of *Mef2c,* for correct L2/3 CPN contralateral S1 axon targeting. These results highlight the multifunctional roles served by Mef2c during cortical development, and they reveal a novel role for Ephrin-Eph signaling, downstream of Mef2c, in controlling homotopic target innervation during the development of intracortical connectivity.

## RESULTS

### Mef2c in post-mitotic neurons directs the development of cortical layers 4 and 5 during embryonic development

An earlier study, employing *Mef2c* deletion starting in neural progenitors (*Nestin-Cre; Mef2c^FL/-^*), identified a role for *Mef2c* in embryonic cortical neuron differentiation and laminar organization^21^. However, it remains unclear in which cell populations Mef2c exerts this early embryonic function, and whether it is important for the development of specific laminar subtypes of cortical neurons.

We first set out to identify the cell-types that express *Mef2c* in the developing murine cortex. Hybridization Chain Reaction (HCR) *in situ* hybridization revealed that *Mef2c* expression is confined to a thin band in the superficial portion of the cortical plate at embryonic day 13.5 (E13.5, Figures 1A and 1A’). At E15.5, *Mef2c* is more broadly expressed throughout the cortical plate (Figures 1B and 1B’). Of note, we did not detect appreciable levels of *Mef2c* in the ventricular and sub-ventricular zones, indicating the *Mef2c* expression is absent in the neural progenitors that populate these regions and that it is restricted to post-mitotic neurons in the cortical plate.

**Figure 1:**
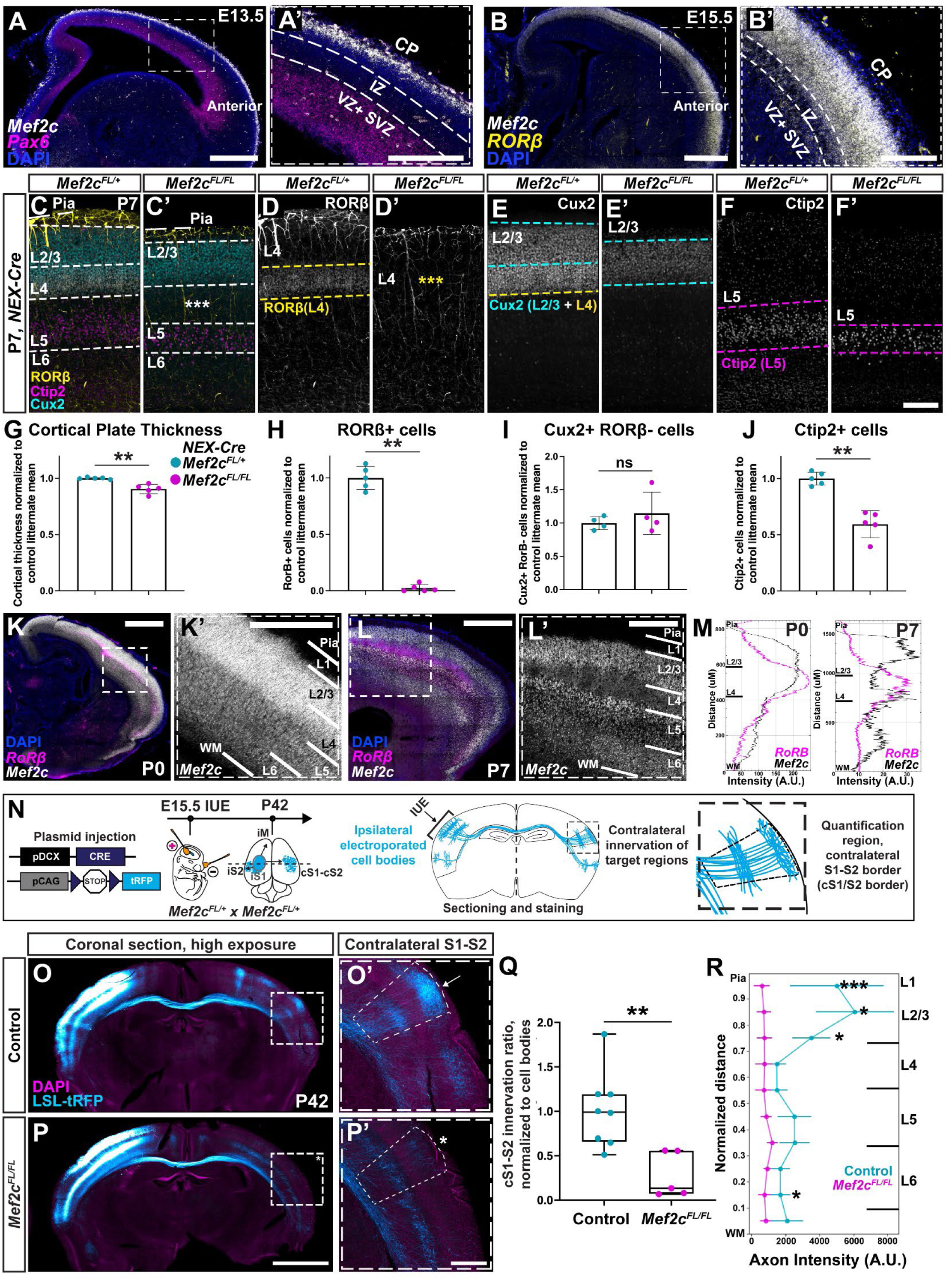
*Mef2c* has dual roles in embryonic cortical laminar organization and postnatal callosal axon projection targeting. **(A and B):** *Mef2c* mRNA expression in sagittal sections at embryonic day 13.5 (E13.5) (A) and E15.5 (B) is limited to post-mitotic neurons in the cortical plate. Insets in A’ and B’: *Mef2c* expression in the cortical plate (CP) and underlying intermediate, sub-ventricular and ventricular zones (IZ, SVZ+VZ). *Pax6* mRNA (magenta in A, A’) marks VZ progenitors and *Rorβ* (yellow in B, B’) marks post-mitotic cortical layer 4 (L4) neurons. **(C-F):** Coronal sections through S1 of postnatal day (P) 7 *Nex-Cre; Mef2c^FL/+^* Control (C, D, E, F) and *Nex-Cre; Mef2c^FL/FL^* mutant (C’, D’, E’, F’) littermates, immunostained for laminar subtype markers. Rorβ+ L4 neurons were absent in the mutant cortex (yellow asterixis in D’), and Ctip2+ L5 neurons were reduced (F and F’), while Cux2+, Rorβ- L2/3 neurons were unaffected (C, E and C’, E’). **(G-J)** Cortical thickness (G) and the number of Layer 5 (L5) Ctip2+ neurons (J) were reduced, and Layer 4 (L4) Rorβ+ neurons were absent (H), while the Layer 2/3 (L2/3) Cux2+, Rorβ- neuron number was unchanged (I) in P7 *Nex-Cre; Mef2c^FL/FL^* mice compared to *Nex-Cre; Mef2c^FL/+^* littermates. Graphs represent mean +/- standard deviation (s.d), n=5 (C, D, F) mice, 3 litters or n=4 (E) mice, 2 litters, for each condition. 3-4 sections per animal were analyzed. Values were normalized to *Nex-Cre; Mef2c^FL/+^* littermate means. Mann-Whitney test, p=0.0079 (G) 0.0079 (H), 0.8857 (I), 0.0079 (J). **(K** and **L):** *In situ* hybridization of (P0) (K) and P7 (L) sagittal sections, and insets (K’, L’) of the S1 barrel field, reveal high *Mef2c* mRNA expression in L2/3 neurons. *Rorβ* (magenta) marks L4 in sensory cortex. **(M)** Fluorescence intensity profiles of *Mef2c* and *Rorβ* expression along the Pia-to-white matter (WM) axis of S1 cortex at P0 (left) and P7 (right), showing high expression of *Mef2c* in both L2/3 and L4 at P0, and a shift to L2/3-specific enrichment at P7. **(N)** *In utero* electroporation (IUE) strategy in a single hemisphere with depicted plasmids for conditional *Mef2c* knockout in post-mitotic S1-L2/3 neurons. Schematics (right) illustrate expected labeling of electroporated cell bodies in S1 cortex and their projections to their major targets: ipsilateral motor cortex (iM), ipsilateral secondary somatosensory cortex (iS2) and the contralateral S1-S2 border (cS1-S2). **(O** and **P)** Coronal sections of electroporated brains representative of *Mef2c^+/+^* and *Mef2c^FL/+^* mice (Control: O, O’) and homozygous *Mef2c^FL/FL^* littermates (P, P’) at P42. *Mef2c^FL/FL^* mice display reduced innervation (asterisk) of cS1-S2 (P’), compared to Control (arrow, O’). **(Q)** Quantification of cS1-S2 innervation reveals a reduction in targeting by *Mef2c^FL/FL^* mutant S1-L2/3 neurons. Innervation was measured as integrated tRFP fluorescence in the boxed cS1-S2 region of high-exposure images, divided by electroporated ipsi-L2/3 cell body intensity from low-exposure images for normalization (see Methods). Data are presented as median +/- intra-quartile range (IQR); whiskers represent range. n=8 control and 5 *Mef2c^Fl/Fl^* mice from 3 independent litters. Mann-Whitney U-test, p=0.0062. **(R)** Fluorescence intensity profiles of innervation along the Pia-to-WM axis of the cS1-S2 target domain at P42 show reduced innervation of *Mef2c^FL/FL^* axons throughout the cortical wall. Data presented as mean +/- s.d of innervation intensity in bins that correspond to 1/10^th^ of the normalized Pia-to-WM distance. n=8 control and 5 *Mef2c^Fl/Fl^* mice from 3 independent litters. Mann Whitney U-test with Bonferroni correction for multiple comparisons, p=0.0155 at 0.75 and 0.85 Pia-to-WM; Unpaired T-test with Bonferroni correction, p=0.002 (0.95 Pia-to-WM), p=0.0336 (0.15 Pia-to-WM). ns, not significant; *, p<0.05; **, p<0.01; ***, p<0.005 Scale bars: 500 μm (A, B, K’, L’, P’), 200 μm (A’, B’, F’), 1000 μm (K, L), 2000 μm (P), See also Figure S1 and Figure S2.

In the light of this expression pattern, we asked whether cortical organization deficits are recapitulated upon post-mitotic cortical neuron-specific *Mef2c* deletion. We chose the *Nex-Cre (Neurod6-Cre)*^61^ transgenic line to recombine a conditional *Mef2c-floxed* (*Mef2c^FL^*) allele, in which the second coding exon is flanked by loxP sites^62^, in all post-mitotic neurons of the dorsal telencephalon starting at E11.5.

We assessed laminar organization in the primary somatosensory cortex (S1) at postnatal day 7 (P7) and observed a complete loss of Rorβ+(Rorb+) Layer 4 (L4) neurons in *Nex-Cre; Mef2c^FL/FL^* animals (Figures 1D, 1D’ and 1H). This uncovers a novel role for Mef2c in the development of cortical L4. The number of Ctip2+ (Bcl11b+) Layer 5 (L5) neurons was also reduced in *Nex-Cre; Mef2c^FL/FL^* animals (Figures 1F, 1F’ and 1J), as was cortical thickness (Figures 1C, 1C’ and 1G), recapitulating observations from the *Nestin-Cre; Mef2c^FL/-^* condition^21^. The number and laminar position of L2/3 neurons was unchanged between control and mutants (Figures 1N, 1N’ and 1R). Differences in Layer 6 (Tle4+) were also not observed (data not shown). Rorβ+ L4 neurons are the main recipients of thalamic input in primary sensory cortical areas and are innervated by Vglut2+ thalamocortical axons, which segregate into barrel-like structures in S1 (Figures S1P and S1Q). However, they fail to do so in *Nex-Cre; Mef2c^FL/FL^* animals that lack L4 (Figures S1P’ and S1Q’).

We next assessed cortical laminar organization in *Nex-Cre; Mef2c^FL/FL^* brains at E16.5, when L4 neurons have migrated into the cortex. The expression of *Rorβ*, a key determinant of L4 neuron identity in sensory cortex^63,64^, also increases in the cortical plate (CP) at this stage^65^. The CP *Rorβ* expression domain was greatly reduced in *Nex-Cre; Mef2c^FL/FL^* animals compared to *Nex-Cre; Mef2c^FL/+^* littermates at E16.5 (Figures S1A-S1B’). Specifically in the somatosensory region, *Rorβ* signal intensity was significantly lower in the *Nex-Cre; Mef2c^FL/FL^* mutants compared to heterozygous littermates (Figures S1D vs 1D’, and 1H). The number of Brn2+ (Pou3f2+) neurons (Figures S1E, S1E’ and S1J) and Ctip2+ neurons (Figures S1F, S1F’ and S1K) in the cortical plate was not significantly different between these two conditions. CP thickness (Figures S1C, S1C’ and S1G) and the total number of nuclei in the CP (DAPI+, Figure S1I) were also unaltered. Reduced *Rorβ* expression coupled with normal expression of other laminar subtype markers and cell numbers in the cortical plate supports *Mef2c* specifically controlling acquisition of L4 neuron identity.

Since *Mef2c* has been reported to impact neuronal survival^26,27^, we asked whether the observed laminar subtype losses are due to elevated cell death. We observed similar levels of cleaved-Caspase 3 (CC3) staining around the midline, a site of normal apoptosis, in both controls and mutants (Figures S1M, S1M’, S1O and S1O’), validating our approach to detect cell death. However, we did not observe elevated CC3 in *Nex-Cre; Mef2c^FL/FL^* S1 cortex, compared to controls, at either E16.5 or P0 (Figures S1L, S1L’, S1N and S1N’). This result rules out increased apoptotic vulnerability of S1-L4 neurons in the absence of *Mef2c* and instead supports mis-specification to a different identity that is not marked by commonly employed cortical laminar subtype markers.

Taken together, our results localize *Mef2c* function to post-mitotic neurons during embryogenesis, and they reveal a new role for *Mef2c* in the development of specific cortical neuron laminar subtypes, namely L4 and L5.

### Mef2c regulates L2/3 callosal projection neuron axon targeting during postnatal development

Next, we sought to determine whether, in addition to its role in laminar organization during embryogenesis, Mef2c also contributes to postnatal cortical development. As a first step, we documented *Mef2c* expression during the first postnatal week. We observed strong *Mef2c* expression in superficial cortical layers (L2/3 and L4) of S1 at P0 (Figures 1K, 1K’ and 1M left), which then shifts to a pattern of laminar-specific enrichment in L2/3 at the end of the first postnatal week (Figures 1L, 1L’ and 1M right). This shift from earlier broad embryonic *Mef2c* cortical expression to robust L2/3-specific expression during the first postnatal week has been previously documented^40^, but its functional relevance is unknown. Since this developmental period coincides with the elaboration of axonal projections by L2/3 callosal projection neurons (L2/3 CPNs), we hypothesized that *Mef2c* plays a role in axon outgrowth and/or guidance.

Pan-cortical deletion of *Mef2c* disrupts general organization of the cortex and is hence unsuitable for assessing cell-autonomous contributions of *Mef2c* to L2/3 axon projection. Therefore, we employed *in utero* electroporation (IUE) of a plasmid encoding Cre recombinase (*pDcx-Cre*, restricted to post-mitotic neurons by the *Doublecortin* promoter^66^) into *Mef2c^Floxed^* embryos at E15.5 to selectively target L2/3 CPNs (Figure 1D). A Cre-dependent fluorescent reporter plasmid (*pCAG-LSL-turboRFP*^67^ or *pCAG-LSL-EGFP*^68^) was co-electroporated to visualize neuronal morphology. We observed normal cortical laminar organization, including L4, upon E15.5 Cre-IUE into *Mef2c^FL/FL^* embryos (Figure S2A), validating the suitability of this approach for L2/3 CPN-autonomous perturbation.

In control *Mef2c^+/+^*and *Mef2c^FL/+^* electroporated animals, S1-L2/3 CPNs send axon projections across the corpus callosum to generate a prominent innervation column at the boundary of contralateral S1 and secondary somatosensory cortex (S2) (Figures 1O and 1O’). However, we observed a strong reduction in homotopic innervation of this contralateral cortical domain upon Cre-IUE into S1-L2/3 CPNs in *Mef2c^FL/FL^* animals (Figures 1P and 1P’). Quantification of axonal fluorescence at the contralateral S1-S2 (cS1-S2) border, normalized to ipsilateral L2/3 electroporation intensity (see Methods), was significantly lower in mutant brains compared to pooled *Mef2c^+/+^*and *Mef2c^FL/+^* littermate controls (Figure 1Q). Control and mutant electroporated brains showed similar levels of ipsilateral L2/3 cell labeling (Figures S2B, S2B’ and S2C), indicating an innervation deficit independent of electroporation efficiency. Intensity profiles of innervation, along the Pia-to-WM axis, revealed a reduction in both deep and superficial layers of the cS1-S2 domain in *Mef2c^FL/FL^* mutants compared to controls (Figure 1R). We also observed reduced innervation of domains more lateral to cS1-S2 in *Mef2c^FL/FL^* mutants compared to controls (Figures S2D-S2E’). Reduced cS1-S2 innervation was not compensated by increased innervation of more medial domains within the barrel field (Figures S2F-S2H).

S1-L2/3 neurons from *Mef2c^FL/FL^* brains also displayed laminar specific innervation deficits at ipsilateral long-range targets. Innervation was reduced in superficial, but not deep, ipsilateral S2 layers (Figures S2K-S2L’) and also in motor cortex (Figure S2M and S2N). Additionally, we observed ectopic subcortical projections to the basolateral amygdala in *Mef2c^FL/FL^* brains (Figures S2K’’ and S2L’’).

These results reveal that *Mef2c* is crucial for targeting S1-L2/3 CPN axons to homotopic domains in the contralateral cortex, and that it is also essential for guiding innervation of superficial, but not deep, layers of distal ipsilateral targets.

### Dual labeling of WT and *Mef2c* conditional-mutant axons in the same brain confirms cell autonomy of Mef2c function for callosal projection targeting

Our IUE-based L2/3-specific *Mef2c* loss-of-function (LOF) paradigm strongly supports a cell autonomous role for Mef2c in regulating homotopic targeting of callosal projections. We further confirmed cell autonomy by comparing target innervation by WT and *Mef2c* conditional-mutant callosal axons in the same brains by employing IUE-based dual-labeling (Figure 2A). We co-electroporated a lower concentration of the *pTRE-Cre* with higher levels of the *pCβA-Flex* construct ^69^, which expresses *tdTomato* in the absence of Cre-recombination. In this scenario only a fraction of *pCβA-Flex* neurons also express *pTRE-Cre*^67^, and in those neurons *tdTomato* is excised and instead EGFP is expressed. To ensure maximum recombination and robust GFP-labeling, we also co-electroporated the Cre-dependent *pCAG-LSL-EGFP-ires-tTA*^67^; tTA positively feeds back to maximize *Cre* expression from the *TRE* promoter. As expected (Figure 2B), we observed nearly exclusive labeling of electroporated L2/3 neurons with EGFP (Cre+) and tdTomato (Cre-), and only a small fraction of double labeling (Figure S3A).

**Figure 2:**
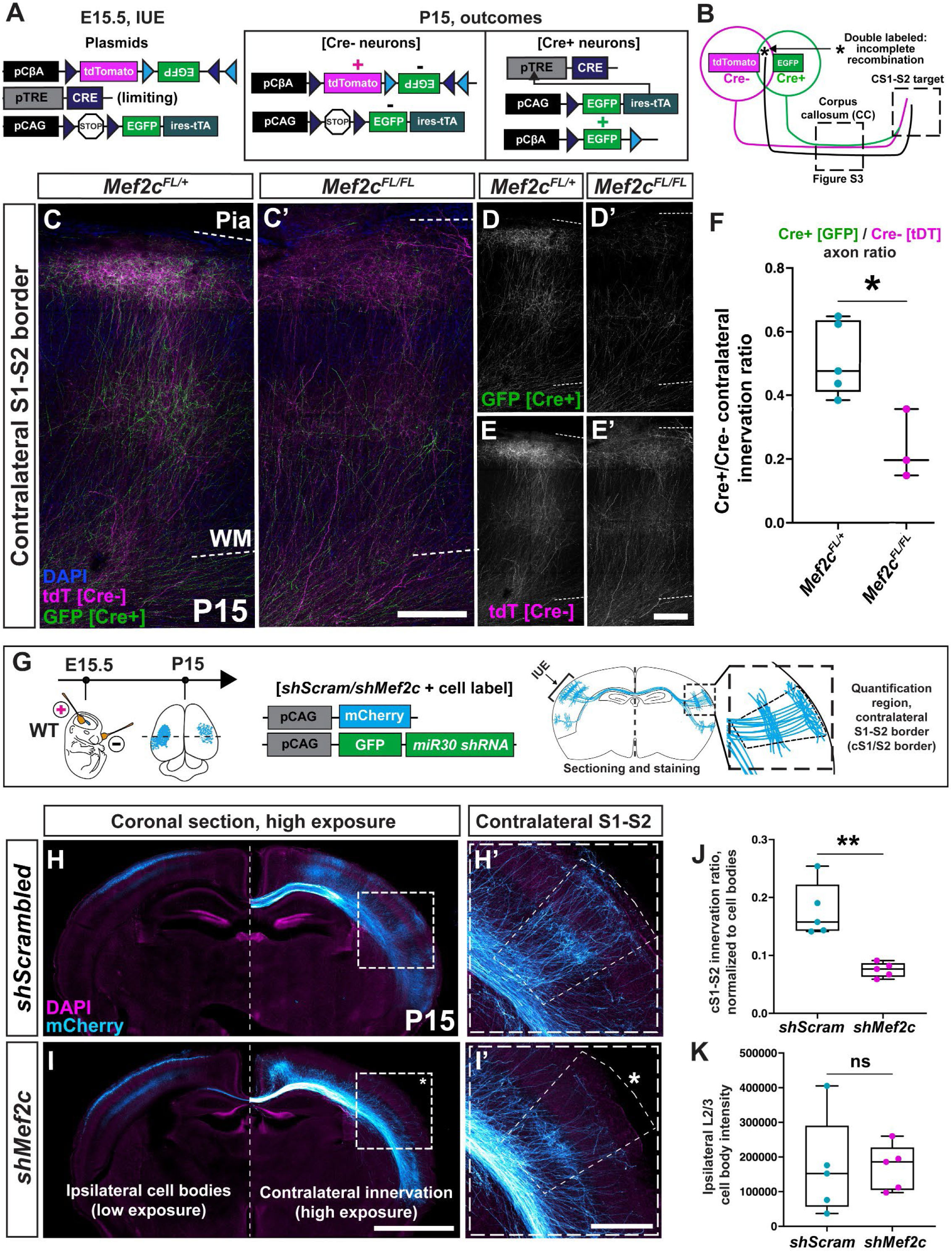
Dual labeling of Cre+ and Cre- callosal projections in *Mef2c-cKO* brains, and L2/3 CPN specific *Mef2c* knockdown independently confirm cell autonomy of *Mef2c* function in CPN axon targeting. (A) Schematic of IUE-based dual-color labeling strategy to mark Cre+ neurons with EGFP and Cre- neurons with tdTomato in the same brain. **(B)** Schematic representation of dual-labeling IUE outcome. All Cre+ neurons express EGFP, but a subset can co-express tdTomato due to incomplete recombination and/or perdurance of tdTomato. **(C-E)** A strong reduction of EGFP (Cre+) innervation was observed at cS1-S2 in the *Mef2c^FL/FL^* brain (C’, D’) compared to tdTomato (Cre-) innervation in the same brain (C’, E’). In the *Mef2c^FL/+^* littermates, both EGFP and tdTomato innervation of cS1-S2 are comparable (C–E). **(F)** Quantification of innervation by Cre+ axons in cS1-S2 to Cre- axons (ratio). *Mef2c^FL/FL^* brains display a significantly lower ratio of Cre+: Cre- axons when compared to *Mef2c^FL/+^* control littermates. Data are presented as median +/- IQR; whiskers represent range. n=5 *Mef2c^FL/+^* and 3 *Mef2c^FL/FL^* mice from one litter. Mann-Whitney U- test, p=0.0357. **(G)** Schematic of IUE-based *Mef2c* knockdown using shRNA plasmids combined with an mCherry cell fill. **(H** and **I)** WT mice electroporated at E15.5 with shRNAs in S1. S1-L2/3 CPNs show reduced innervation of cS1-S2 upon *Mef2c* shRNA IUE (asterisk, I’), compared to littermates electroporated with Scrambled Control shRNA (H, H’). **(J** and **K)** Quantification of cS1-S2 innervation, normalized to ipsilateral cell body intensity, reveals reduced cS1-S2 innervation by *shMef2c* neurons compared to *shScrambled* controls (J). IUE efficiency, measured by ipsilateral cell body intensity, was not significantly different between mutant and control. (Q). Data are presented as median +/- IQR; whiskers representing the range. n=5 *shScrambled* and 5 *shMef2c* mice from 2 independent litters. p=0.0079 (P) and 0.5476 (Q) Mann-Whitney U-test. ns, not significant; *, p<0.05; **, p<0.01. Scale bars: 200 μm (C’, E’), 2000 μm (I), 500 μm (I’) See also Figure S3.

Both Cre+ and Cre-neurons innervated the cS1-S2 target in controls (Figures 2C, 2D and 2E), however, only Cre-, but not Cre+, neurons innervated cS1-S2 in *Mef2c^FL/FL^* brains (Figures 2C’, D’ and E’). Quantification of the ratio of Cre+:Cre- innervation of cS1-S2 revealed a reduction in mutants compared to controls (Figure 2F). Cre+:Cre- intensity ratio at the corpus callosum midline was however not significantly different between the two genotypes, indicating no deficit in midline crossing (Figures S3B-D). By incorporating controls within the same brain, these results confirm the cell autonomy of *Mef2c* contralateral S1-L2/3 CPN target innervation defects.

The *Mef2c^Floxed^* allele produces a deletion of a large portion of the Mef2c DNA-binding domain, but since the floxed exon is in-frame the rest of the protein is unaltered. We therefore employed shRNA-based knockdown of *Mef2c,* by IUE in L2/3 neurons, as an additional method for assessing *Mef2c* LOF. Contralateral innervation deficits were recapitulated in this paradigm of *Me2c* LOF (Figures 2G-2K, validation of knockdown in Figures S3E-S3F’’), further demonstrating that Mef2c function is crucial for callosal axon target innervation.

### *Mef2c* specifically regulates target innervation, and not midline crossing, of callosal projection neurons

L2/3 CPN midline crossing appears unaffected in *Mef2c^FL/FL^* brains at both P42 (Figures S2O and S2P) and P15 (Figures S3B-D), suggesting a specific defect in the later process of axon targeting. Ectopic projections to heterotopic cingulate cortical domains in *Mef2c^FL/FL^* brains (Figures S2I and S2J) further strengthen the case for a defect in axon targeting rather than in outgrowth or midline-crossing. We sought to confirm the specific function of *Mef2c* in callosal projection targeting, and not midline crossing, by comparing the time-course of *Mef2c^FL/+^* and *Mef2c^FL/FL^* mutant S1-L2/3 callosal projection development.

At P4 and P8, we observed normal midline crossing in both conditions (Figures 3A-3F), confirming that *Mef2c* is dispensable for midline crossing. At P8, control S1-L2/3 CPN axons broadly innervate the entire contralateral S1 cortical domain (Figures 3C and 3G). Mutant axons are confined to the white matter and do not enter the cortical plate (Figures 3D, 3H and 3I), confirming that *Mef2c* is required for the initial target innervation of S1-L2/3 CPN axons into the homotopic contralateral domain. During the second postnatal week, control S1-L2/3 CPN axons in the more medial contralateral S1 are eliminated, while those at the cS1-S2 border persist and elaborate horizontal branches in L2/3 and L5 to produce the columnar innervation characteristic of more mature stages (Figures 3J and 3J’). *Mef2c^FL/FL^* mutant axons remain confined to the white matter (Figures 3K-L), ruling out the possibility that target innervation is recovered after a delay.

**Figure 3:**
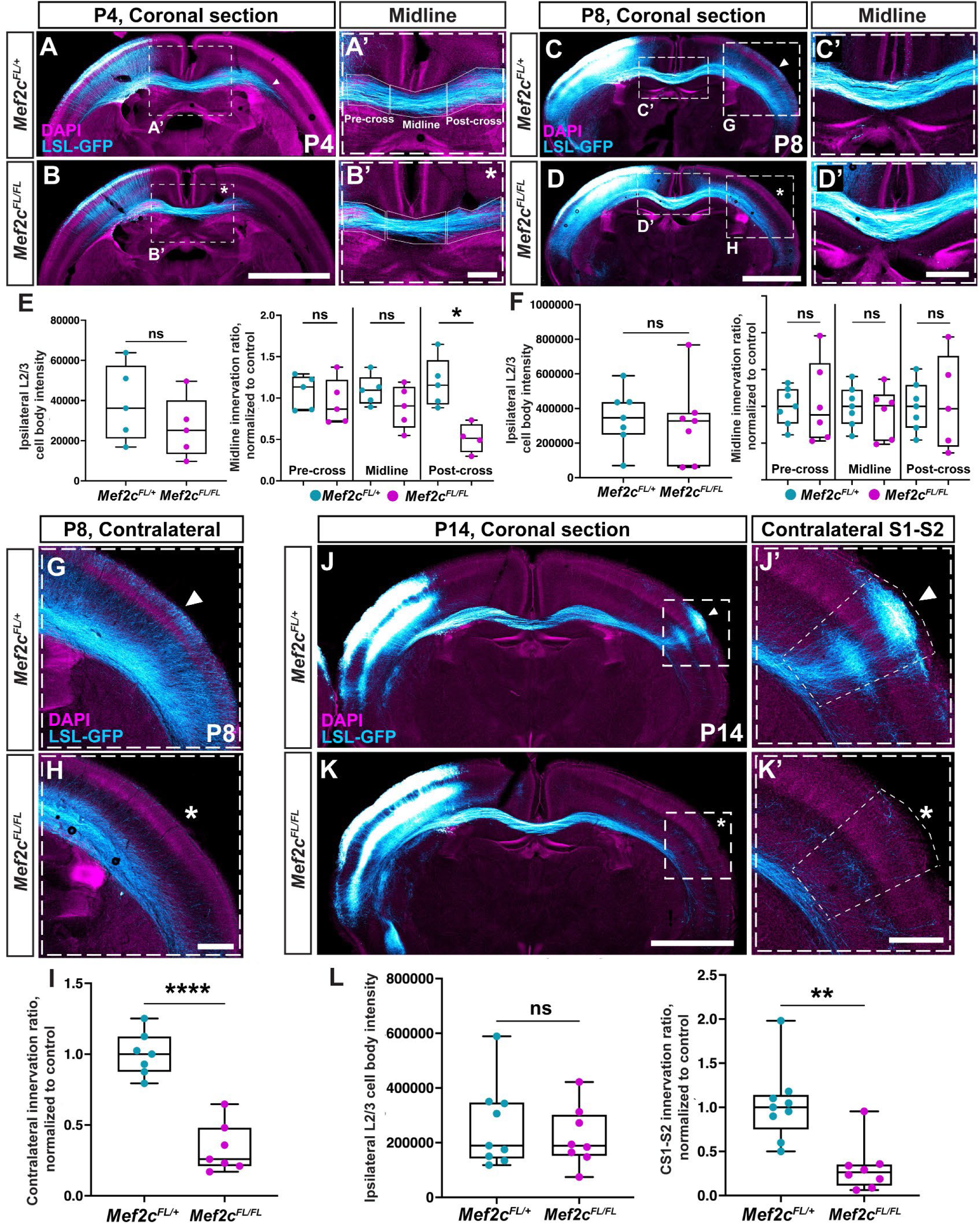
*Mef2c* specifically regulates target innervation, and not midline crossing, of callosal projection neurons. **(A** and **B)** Both *Mef2c^FL/+^* (A, A’) and *Mef2c^FL/FL^* (B, B’) E15.5-Cre-IUE brains exhibit normal midline crossing of S1-L2/3 CPNs at P4. However, post-crossing axon outgrowth is impaired for *Mef2c^FL/FL^* axons (asterisks in B, B’) compared to *Mef2c^FL/+^* littermates (arrowhead, A). **(C** and **D)** *Mef2c^FL/+^* control (C, C’) or *Mef2c^FL/FL^*(D, D’) E15.5-Cre-IUE brains display no deficits in midline crossing or post-crossing white matter extension at P8. **(E)** IUE efficiency, measured as ipsilateral-cell body fluorescence, was not significantly different between the *Mef2c^FL/+^* and *Mef2c^FL/FL^* analyzed at P4. Signal intensity in the pre-crossing and midline regions of the corpus callosum, normalized to IUE-efficiency (see Methods), were not significantly different between *Mef2c^FL/+^* and *Mef2c^FL/FL^;* while there was a significant reduction in post-crossing corpus callosum signal intensity from *Mef2c^Fl/Fl^* S1-L2/3 CPNs at P4 (right). Data are presented as median +/- IQR; whiskers represent range. n=5 *Mef2c^FL/+^* and 5 *Mef2c^FL/FL^* mice, 2 litters. Mann-Whitney U-test p=0.3095 (cell bodies), p=0.5476 (pre-cross), p=0.3095 (midline), p=0.0159 (post-cross). **(F)** IUE efficiency (left) and signal intensity in the pre-crossing, midline and post-crossing of the corpus callosum, normalized IUE-efficiency (right), were not significantly different between *Mef2c^FL/+^* and *Mef2c^FL/FL^* brains at P8, indicating recovery of the initial axon extension deficit seen at P4. Data are presented as median +/- IQR; whiskers represent the range. n=7 *Mef2c^Fl/+^* and 7 *Mef2c^Fl/Fl^* mice, 2 litters. Mann-Whitney U-test: p=0.5350 (cell bodies), p=0.8357 (pre-cross), p=0.6282 (midline), p=0.8767 (post-cross). **(G** and **H)** Higher magnification images of the contralateral cortex at P8, showing that *Mef2c^FL/+^* S1-L2/3 axons invaded the contralateral cortical plate (arrowhead in G), while *Mef2c^FL/FL^* mutant S1-L2/3 axons were restricted to the white matter (asterisk in H). **(I)** Quantification of total innervation in the contralateral S1 cortical plate, normalized to ipsilateral L2/3 cell body intensity, showed a significant reduction in innervation by *Mef2c^FL/FL^* S1-L2/3 neurons compared to *Mef2c^FL/+^* controls. Data are presented as median +/- IQR; whiskers represent range. n=7 *Mef2c^Fl/+^* and 7 *Mef2c^FL/FL^* mice, 2 litters. Mann-Whitney test: p=0.0006. **(J** and **K)** *Mef2c^FL/+^* (J, J’) or *Mef2c^FL/FL^* (K, K’) E15.5-Cre-IUE brains assessed at P14. *Mef2c^FL/+^* S1-L2/3 CPN innervation is refined into columns targeting specific domains, like the cS1-S2 border (J’). Mutant S1-L2/3 axons remained confined to the contralateral white matter (K, K’), and failed to innervate cS1-S2 (asterisk, K’). **(L)** Quantification of ipsilateral L2/3 cell body fluorescence (left) showed no significant difference between *Mef2c^FL/+^* and *Mef2c^FL/FL^* brains at P14(left). Quantification of cS1-S2, innervation, normalized to ipsilateral L2/3 cell body intensity, showed a significant reduction for *Mef2c^FL/FL^* S1-L2/3 neurons compared to *Mef2c^Fl/+^* (right). Data are presented as median +/- IQR; whiskers represent range. n=9 *Mef2c^Fl/+^* and 8 *Mef2c^Fl/Fl^* mice, 3 litters. Mann-Whitney test: p=0.8148 (cell bodies) and p=0.0010 (innervation). *, p<0.05; **, p<0.01; ****, p<0.001 Scale bars: 2000 μm (B, D, K), 500 μm (B’, D’, H, K’)

These results show that *Mef2c* broadly regulates intracortical axon target innervation without affecting axon outgrowth and midline crossing.

### Mef2c mutant L2/3 neurons exhibit aberrant connectivity at intercortical target domains

Is the loss of contralateral target innervation by *Mef2c* mutant S1-L2/3 CPN axons accompanied by a concomitant decline in neuronal connectivity? To answer this question, we co-electroporated a construct encoding the Cre-dependent, mCherry-tagged, monosynaptic WGA (mWmC) anterograde tracer^70^ along with *pDCX-Cre* and *pCAG-LSL-EGFP* (cell label) constructs into control (*Mef2c^+/+^* and *Mef2c^FL/+^*) or *Mef2c^FL/FL^* E15.5 embryos. We then assessed connectivity as revealed by trans-synaptic labeling at P42. In control brains, we detected several neurons with perisomatic mWmC fluorescence in L5 and L2/3 of cS1-S2, coinciding with GFP+ axonal innervation (Figures 4C-4C’ and 4E-4F’), strongly suggesting that they were synaptically connected to electroporated S1-L2/3 CPNs in the ipsilateral cortex. In *Mef2c^FL/FL^* mutant brains, the number of connected mWmC+ neurons in cS1-S2 was greatly reduced in both L2/3 and L5 (Figures 4D-4D’, 4G-4H’ and 4J), reflecting the reduced axon innervation we observed in the absence of Mef2c (Figure 4I).

**Figure 4:**
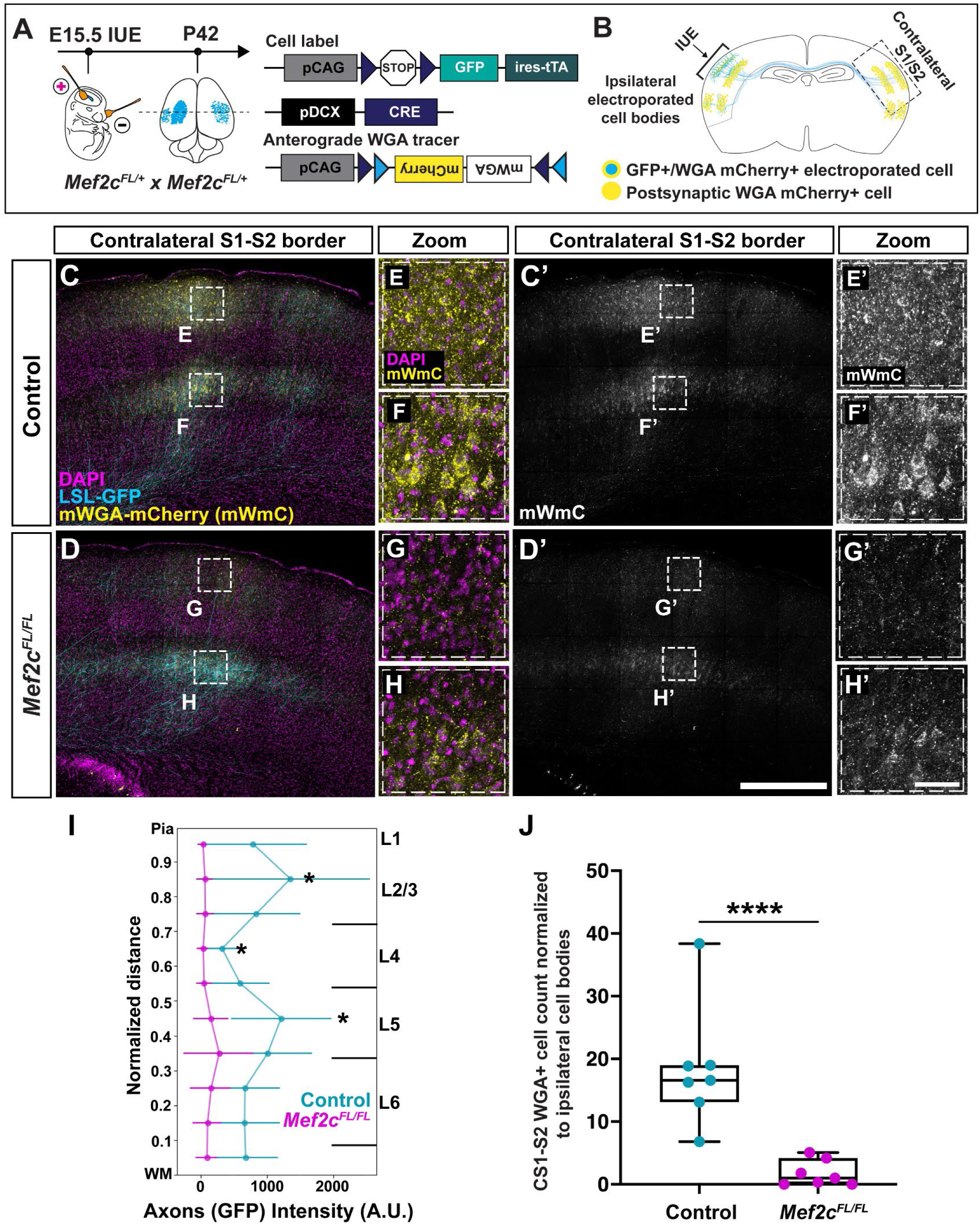
WGA anterograde tracing suggests *Mef2c* mutant S1-L2/3 CPNs fail to form synapses with contralateral cortical targets. **(A)** IUE experiment workflow to introduce plasmids expressing Cre, the Cre-dependent anterograde synaptic tracer mWGA-mCherry (mWmC), and a cell-label (GFP) in *Mef2c^FL/FL^* embryos. **(B)** Schematic representation of IUE outcome. Electroporated neurons in ipsilateral S1 are double-labeled with GFP and mCherry. At contralateral target regions, only neurons post-synaptic to the electroporated neurons will express perisomatic mCherry. **(C** and **D)** Images of cS1-S2 reveal a reduction in both GFP axonal innervation and mWmC post-synaptic partners of S1-L2/3 CPNs in the *Mef2c^FL/FL^* brain (D, D’) compared to those from Control littermates (C, C’). **(E-H)** Multiple neurons show perisomatic mWmC as can be seen in high magnification insets of L2/3 (E) and L5 (F) in the Control brain, while almost none are detected in the *Mef2c^FL/FL^* brain (G and H). **(I)** Fluorescence intensity profiles of axon innervation along the Pia-to-WM axis in cS1-S2 show reduced innervation of *Mef2c^FL/FL^* axons throughout the cortical wall. Data presented as mean +/- s.d of innervation intensity in bins that correspond to 1/10^th^ of the normalized Pia-to-WM distance. Mann Whitney U-test with Bonferroni correction for multiple comparisons: p=0.0233 at 0.85, 0.65 and 0.45 Pia-to-WM. **(J)** Quantification of mWmC^+^ cell-body number in cS1/S2 reveals *Mef2c^FL/FL^* S1-L2/3 neurons appear to synapse onto fewer targets compared to littermate Controls. Cell counts from L2/3 and L5 in the cS1-S2 were summed and divided by ipsilateral L2/3 GFP intensity (from separate low-exposure images, see Methods) for normalization. Mann-Whitney U-test, p=0.0006. n=7 Control and 7 *Mef2c^Fl/Fl^* animals from 2 litters; *, p<0.05; ****, p<0.001. Scale bars: 500 μm (D’), 50 μm (H’). See also Figure S4.

Local connectivity with partners in ipsilateral S1-L2/3 and L5 appeared grossly similar between controls and mutants (Figures S4A-S4F’). At long range targets of S1-L2/3 neurons in the ipsilateral cortex, where innervation of superficial layers is reduced in *Mef2c^FL/FL^* IUE brains, we observed a corresponding decrease in the number of mWmC+ connected neurons in superficial layers (Figures S4G-S4J). In addition, the basolateral amygdala, which is ectopically targeted by *Mef2c^FL/FL^* S1-L2/3 CPN axons, was also densely packed with mWmC+ neurons, indicating that mis-targeted mutant axons appear capable of forming synaptic connections.

Altogether, these results show *Mef2c* mutant L2/3 CPNs exhibit axon targeting deficits reflected in a corresponding loss of connectivity with many contralateral S1 targets.

### *EphA6* expression is upregulated in Mef2c-mutant S1 L2/3 neurons

How might *Mef2c* direct axonal targeting of a specific population of cortical neurons? One possibility is that *Mef2c* is responsible for transcriptional control of L2/3 laminar and/or S1-areal identity, and axonal mistargeting of *Mef2c-*mutant S1-L2/3 CPNs arises due to these neurons adopting a different identity with distinct projection targets. Alternatively, the expression of specific axon guidance genes that regulate callosal projection targeting could be misregulated upon *Mef2c* LOF.

To test the first hypothesis, we evaluated the expression of L2/3 laminar identity markers (Brn2, Figures S5A-S5C and Cux1, Figures S5D-S5F) and the S1 areal marker Bhlhb5/Bhlhe22 (Figures S5G-S5I) in Cre-electroporated *Mef2c^FL/FL^* neurons, compared to *Mef2c^FL/+^* controls. We did not find any differences in marker expression between the two conditions, implying that post-mitotic *Mef2c* is dispensable for broad establishment of L2/3 laminar and S1 areal identities. Nor did we observe elevated cell death in Cre- electroporated *Mef2c^FL/FL^* mutants (Figures S5J-S5L), ruling out projection deficits in the mutant resulting from a loss of electroporated neurons.

Next, we consulted a published dataset comparing the bulk transcriptomes of pan-cortical *Emx-Cre; Mef2c^FL/FL^* mutant P21 cortices with *Mef2c^FL/FL^* controls^34^ to identify candidate cell-surface molecules that might specifically affect callosal projection targeting. The expression of several members of the *EphA* family of repulsive guidance receptors, *EphA5, EphA6 and EphA7,* whose cognate repulsive ligand *EfnA5* is expressed in S1, are all upregulated in *Emx-Cre; Mef2c^FL/FL^* mutants (Figure 5A). Increased *EphA* receptor expression in *Mef2c-*mutant S1-L2/3 CPNs has the potential to sensitize them to repulsion by the cognate ligand EfnA5 in contralateral S1^54,58,71^, contributing to the observed innervation deficit in the absence of Mef2c.

**Figure 5:**
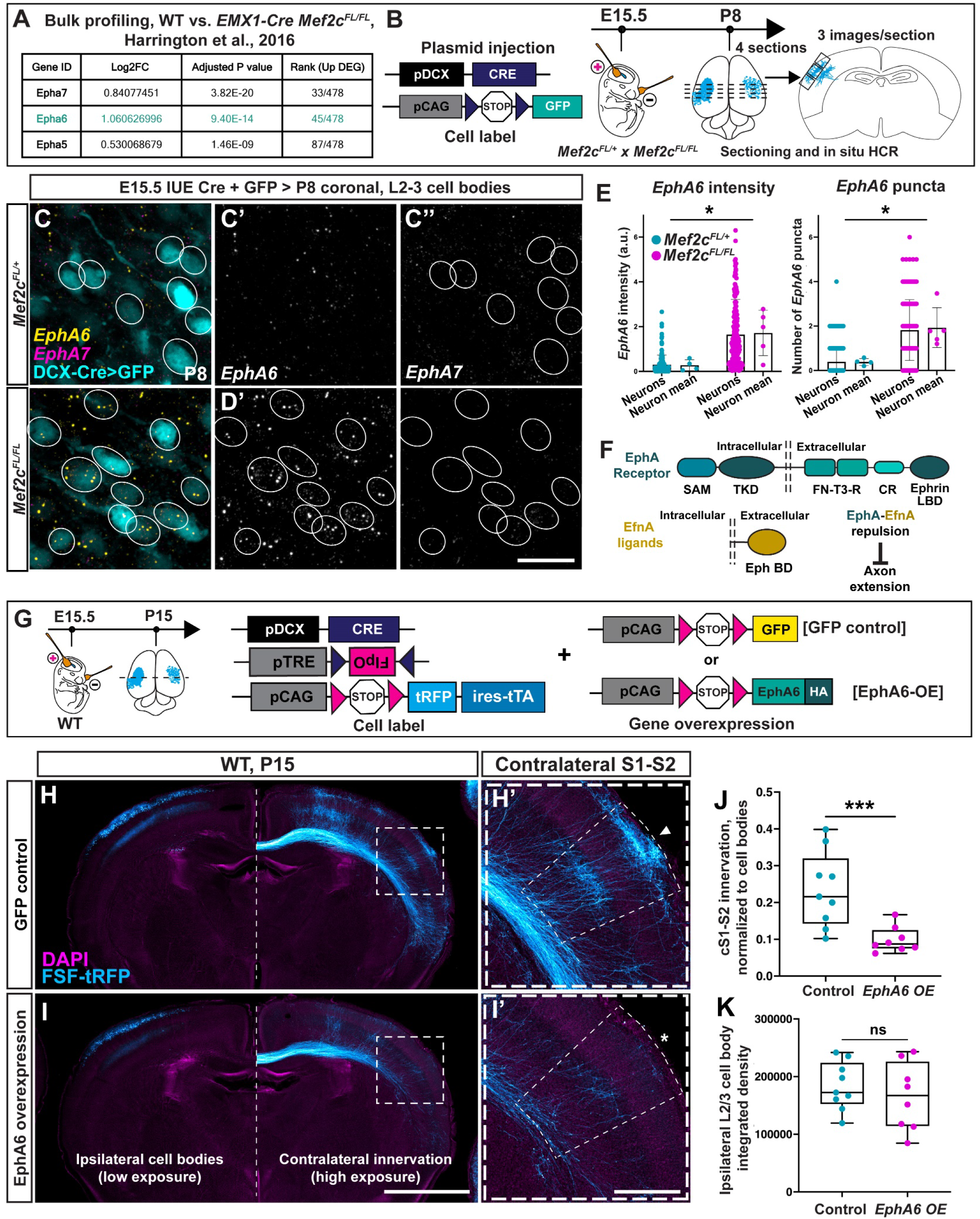
*EphA6* expression is upregulated in *Mef2c-*mutant S1-L2/3 CPNs, and *EphA6* overexpression in WT S1-L2/3 CPNs leads to reduced contralateral target innervation. **(A)** Table details upregulated expression of *Eph* receptors in published bulk transcriptomic profiling of *Emx1-Cre; Mef2c^FL/FL^;* vs *Cre-; Mef2c^FL/FL^* Control P21 cortex^34^. **(B)** Experimental workflow to compare *EphA* receptor expression in Cre-electroporated S1-L2/3 CPNs in *Mef2c^FL/+^* control and *Mef2c^FL/FL^* brains. S1-L2/3 cell bodies were evaluated in 4 different sections from each brain, using 3 fields of view per section. **(C** and **D)** Cre-electroporated P8 S1-L2/3 CPN cell bodies (GFP^+^) in *Mef2c^FL/+^* control (C-C’’) and *Mef2c^Fl/Fl^* (D-D’’) mutant brains, hybridized with probes against *EphA6* and *EphA7*. *EphA6* expression is upregulated in mutant neurons (C’ and D’), while *EphA7* is not differentially expressed (C” and D”). **(E)** Quantification of fluorescence signal intensity (left) and puncta number (right) within GFP+ cell bodies both show a significant upregulation of *EphA6* expression in *Mef2c^FL/FL^* S1-L2/3 neurons. Data are presented as mean +/- s.d for measures from individual neurons (left bars) and mean per neuron for individual mice (right bars) for each genotype. n=145 neurons from N=4 *Mef2c^Fl/+^* control and n=186 neurons from N=5 *Mef2c^Fl/Fl^* mice. Nested T-test: p=0.0276 (intensity), 0.0121 (puncta). **(F)** Domain organization of EphA receptors and Ephrin A (EfnA) ligands. **(G)** Experimental workflow to assess the effect of *EphA6* overexpression, using IUE, on S1-L2/3 neuron contralateral target innervation. **(H** and **I)** *EphA6* overexpression in WT S1-L2/3 neurons reduces innervation of contralateral targets (asterisk, I’) compared to GFP-expressing Control neurons (arrow, H’). **(J** and **K)** Quantification of cS1-S2 innervation, normalized to ipsilateral cell body intensity, shows significant reduction upon *EphA6* overexpression (J). Electroporation efficiency, measured as tRFP signal intensity of ipsilateral L2/3 cell bodies from low-exposure images, was not significantly different between the two conditions (K). Data are presented as median +/- IQR; whiskers represent range; n=9 GFP-control and 8 EphA6-overexpression mice, 3 litters. Mann-Whitney U-test: p=0.0025 (J) and 0.5414 (K). ns, not significant; *, p<0.05; ***, p<0.005. Scale bars: 50 μm (D’’), 2000 μm (I), 500 μm (I’). See also Figure S5.

To validate whether any of these receptors were upregulated specifically in L2/3 CPNs, we performed HCR *in situ* hybridization on P7 brain sections from E15.5-Cre+GFP IUE animals (Figure 5B). We observed that *EphA6* expression, but not *EphA7* expression, is increased in *Mef2c^FL/FL^* S1-L2/3 CPNs, compared to *Mef2c^FL/+^* controls (Figures 5C-5E).

Together, these results support the hypothesis that *Mef2c* regulates callosal projection targeting by controlling the expression of specific guidance molecules, rather than broadly controlling areal and laminar identity acquisition during postnatal development.

### *EphA6* overexpression impairs contralateral target innervation by S1 L2/3 neurons

As a first test of *EphA6-EfnA5* regulation of S1-L2/3 CPN axon contralateral targeting, we assessed the effects of *EphA6* overexpression in WT S1 L2/3 CPNs on axon projection. We first cloned C-terminal HA-tagged EphA6 ORF into a Flp recombinase-dependent expression system developed for gene manipulation in cortical neurons^72,73^. We co-electroporated this construct with *pDcx-Cre*, Cre-dependent Flp (*TRE-DIO-FlpO)*^74^ and *pCAG-FSF-turboRFP-ires-tTA,* a construct which in addition to labeling neurons drives Flp recombinase expression to promote maximum recombination of the Flp dependent constructs, thereby ensuring high expression^68^ (Figures 5G and S5M).

Fluorescence intensity in ipsilateral L2/3, a measure of electroporation efficiency, was comparable between L2/3 CPNs that overexpressed *EphA6* and controls expressing *GFP* (Figures 5H, 5I and 5K). Fluorescence intensity at cS1-S2, normalized to ipsilateral L2/3 fluorescence, was significantly reduced in brains overexpressing *EphA6* compared to *GFP* controls (Figures 5H’, 5I’ and 5J).

These results show that EphA6 overexpression in S1-L2/3 CPNs leads to reduced contralateral target innervation, phenocopying *Mef2c* LOF and lending support to the idea that *EphA6* downregulation by *Mef2c* is important for proper S1 L2/3 CPN axon targeting.

### Loss of EphA6 function partially restores contralateral target innervation in *Mef2c* mutant L2/3 CPNs

If *EphA6* downregulation by Mef2c is crucial for S1-L2/3 CPN contralateral target innervation, *EphA6* LOF is expected to restore target innervation in *Mef2c* mutant S1 L2/3 CPNs. Eph-receptor tyrosine kinase signaling is dependent on cross-autophosphorylation of intracellular domains in dimeric receptors assemblies^75^. To disrupt receptor function, we overexpressed a Flp-dependent *EphA6ΔIntra-Celluar Domain (ICD)-GFP* construct in S1-L2/3 neurons by IUE, again employing our Cre + Flp dependent cell labeling and overexpression strategy (Figure 6A). Receptor constructs lacking the intracellular domains function as dominant-negatives when overexpressed since they are incorporated in dimers with WT receptors, rendering them incapable of signaling^76,77^ (Figure 6B). Overexpression of *EphA6ΔICD-GFP* in *Mef2c^FL/+^* S1-L2/3 CPNs did not alter callosal projection or targeting as compared to expression of *GFP* (Figures S6A-S6C), consistent with EphA6 function being dispensable for these processes.

**Figure 6:**
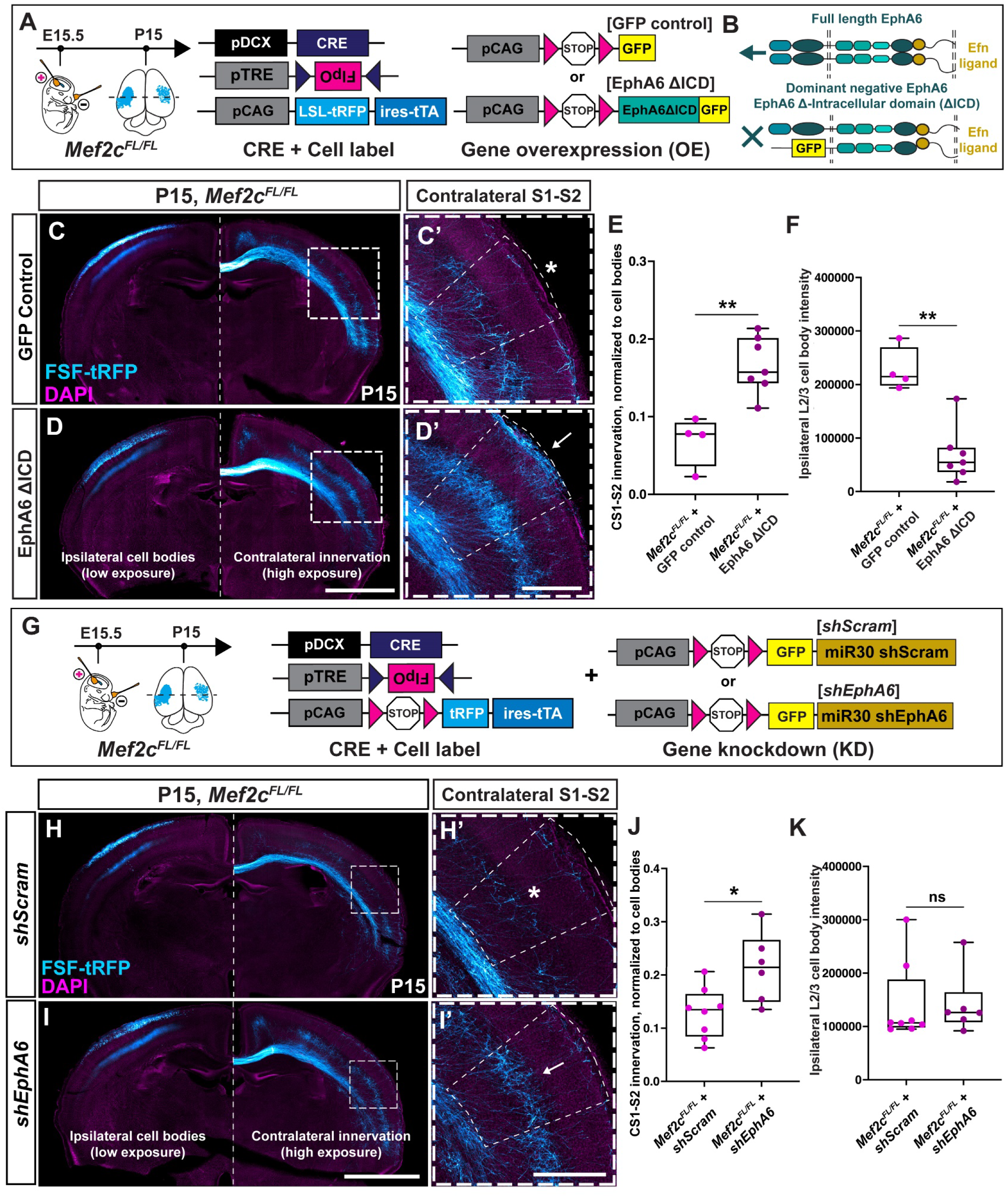
*EphA6* knockdown and dominant-negative expression in *Mef2c* mutant S1-L2/3 CPNs partially restore contralateral axon targeting. **(A)** Experimental workflow to assess the effect of expressing dominant-negative *EphA6ΔICD-GFP* on contralateral target innervation using IUE in *Mef2c^FL/FL^* S1-L2/3 neurons. **(B)** Schematic representation depicting the dominant-negative function of EphA6ΔICD-GFP. **(C** and **D)** *EphA6ΔICD-GFP* expression in *Mef2c^FL/FL^* S1-L2/3 neurons (D) increases contralateral target innervation (arrow, D’) as compared to GFP-expressing *Mef2c^FL/FL^* neurons (C, asterisk in C’). **(E** and **F)** Quantification showing increased cS1-S2 innervation, normalized to ipsilateral cell body intensity, upon *EphA6ΔICD* expression in *Mef2c^FL/FL^* S1-L2/3 neurons compared to *Mef2c^FL/FL^* neurons electroporated with GFP (E). Electroporation efficiency, measured as tRFP signal intensity of ipsilateral L2/3 cell bodies from low-exposure images, was significantly lower for *EphA6ΔICD* IUE compared to GFP IUE in *Mef2c^FL/FL^* brains (F). Data are presented as median +/- IQR; whiskers represent range. n=4 GFP- control and 7 EphA6ΔICD-expressing brains, 3 independent litters. Mann-Whitney U-test: p=0.0061 for both (E) and (F). **(G)** Experimental workflow to assess the effect of EphA6 knockdown, using IUE, on *Mef2c^FL/FL^* S1-L2/3 neuron contralateral target innervation. **(H** and **I)** *EphA6* knockdown in *Mef2c^FL/FL^* S1-L2/3 neurons (I) increased innervation of contralateral targets (arrow, I’) compared to Scrambled shRNA-expressing *Mef2c^FL/FL^* neurons (H, asterisk in H’). **(J** and **K)** cS1-S2 innervation, normalized to ipsilateral cell body intensity, is significantly increased upon *EphA6* knockdown in *Mef2c^FL/FL^* S1-L2/3 neurons compared to *Mef2c^FL/FL^* + scrambled controls (J). Electroporation efficiency, measured as tRFP signal intensity of ipsilateral L2/3 cell bodies from low-exposure images, was not significantly different between the two groups (K). Data are presented as median +/- IQR; whiskers represent range. n=8 *shScram*-control and 6 *shEphA6* brains, 4 litters. Mann-Whitney U-test: p=0.0293 (J) and 0.4136 (K). ns, not significant; *, p<0.05; **, p<0.01. Scale bars: 2000 μm (D, I), 500 μm (D’, I’) See also Figure S6.

*Mef2c^FL/FL^* S1-L2/3 CPNs expressing *EphA6ΔICD-GFP* displayed significantly higher innervation of the cS1-S2 contralateral target compared to *Mef2c^FL/FL^* S1 L2/3 CPNs that expressed only GFP (Figures 6C-6E). We also observed lower ipsilateral L2/3 fluorescence intensity in *Mef2c^FL/FL^ + EphA6ΔICD-GFP* brains compared to *Mef2c^FL/FL^ + GFP* brains (Figure 6F). This is likely due to a difference in IUE efficiency between the two groups in this experiment. However, our quantification of cS1-S2 innervation involves dividing cS1-S2 fluorescence intensity by ipsilateral L2/3 fluorescence intensity to obtain a normalized innervation assessment, which is agnostic to differences in IUE efficiency. Therefore, these results show that disruption of EphA6 receptor signaling in *Mef2c* mutant L2/3 CPNs significantly restores contralateral target innervation.

In addition to disruption of EphA6 signaling using a dominant-negative EphA6 receptor, we also employed shRNA-mediated knockdown of *EphA6* to assess its involvement in S1-L2/3 CPN axon targeting downstream of *Mef2c* (Figure 6G). We co-electroporated *EphA6*-targeting (*shEphA6*) or scrambled-control (*shScram)* shRNAs with an HA-tagged *EphA6* construct and observed a loss of HA-immunofluorescence in *shEphA6* brains, validating their efficiency (Figures S6G-S6H’). *shEphA6* expression in *Mef2c^FL/+^* S1-L2/3 CPNs did not alter callosal projection or targeting as compared to expression of *shScram* (Figures S6D-S6F), further demonstrating that *EphA6* is dispensable for callosal axon development.

Unlike in the *EphA6ΔICD-GFP* experiment, we did not observe significant differences in ipsilateral L2/3 fluorescence intensities in between *Mef2c^FL/FL^ + shEphA6* vs. *Mef2c^FL/FL^ + shScram* (Figure 6K). As observed with *EphA6ΔICD-GFP* expression, *Mef2c^FL/FL^* S1-L2/3 CPNs that expressed *shEphA6* also displayed significantly stronger innervation of the cS1-S2 contralateral target compared to *Mef2c^FL/FL^* S1-L2/3 CPNs that expressed *shScram* (Figures 6H-6J).

Taken together, these results show that disruption of *EphA6* function restores contralateral target innervation by *Mef2c* mutant S1-L2/3 CPNs, supporting Mef2c promotion of S1-L2/3 CPN target innervation through downregulation of *EphA6* expression.

### Depletion of EphrinA5 in the contralateral S1 target domain restores innervation by *Mef2c* mutant S1 L2/3 CPN axons

*EphrinA5 (EfnA5)*, which encodes a repulsive ligand for EphA6, is broadly expressed in S1 at the end of the first postnatal week, but not in more anterior/medial areal domains such as motor and cingulate cortex (Figures 7A-7B’). This area specific *EfnA5* expression coincides with the timing of L2/3 intracortical axon target innervation^78^. We hypothesized that *Mef2c* mutant S1-L2/3 CPN axons are sensitive to repulsion by Ephrin ligands due to elevated *EphA6* expression and hence fail to innervate the *EfnA5*-high S1 target domain, as depicted in Figure 7C. Deletion of *EphA6* in this context likely restores contralateral targeting by suppressing sensitivity to EphrinA5 by *Mef2c* mutant CPN axons. This model also predicts that depletion of EfnA5 in the contralateral S1 target domain should have a similar effect.

**Figure 7:**
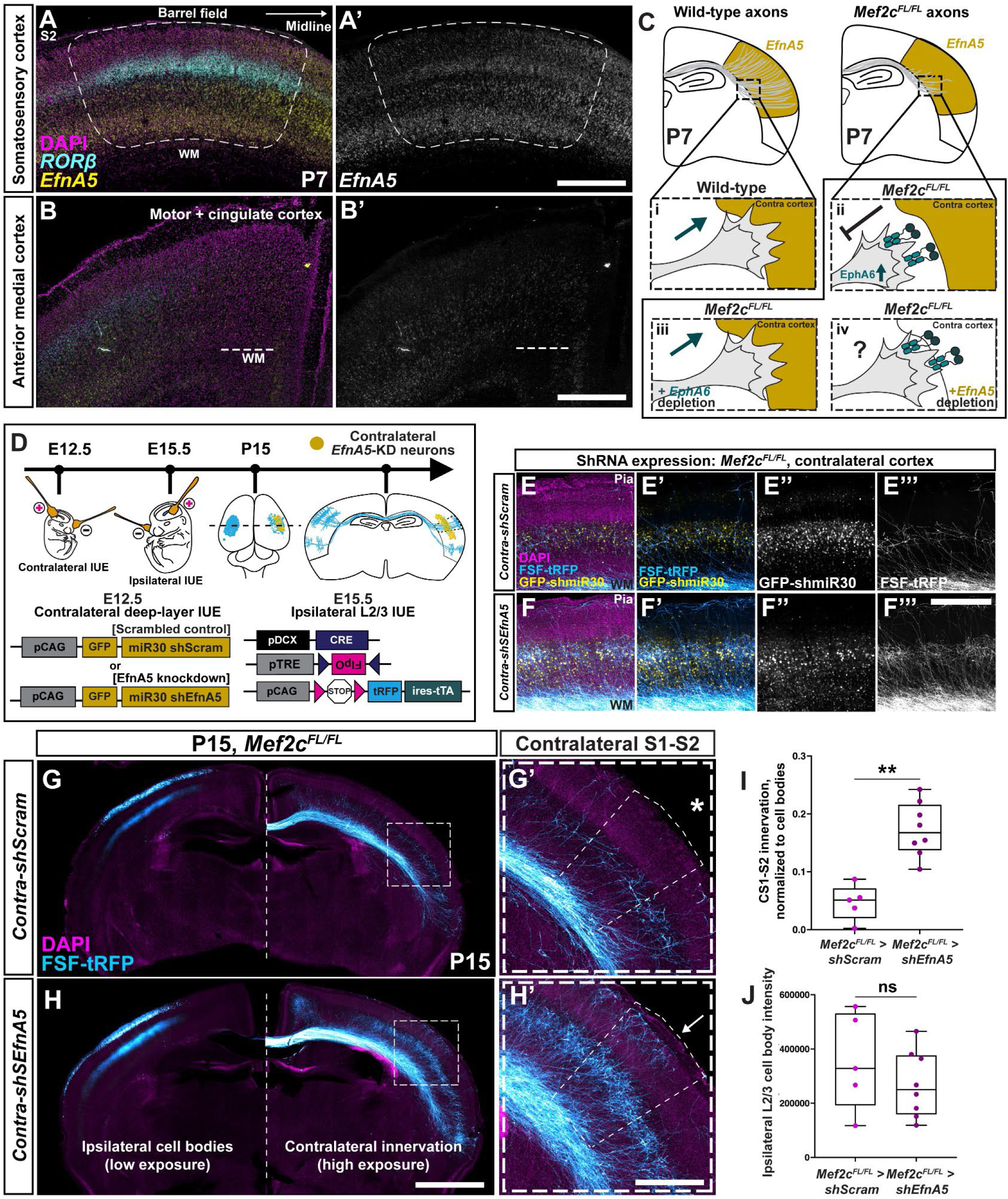
The repulsive ligand *EfnA5* is expressed in the contralateral target domain of S1-L2/3 CPNs, and innervation of *Mef2c* mutant neurons is partially restored upon EfnA5 depletion at the target. **(A)** HCR *in situ* hybridization in coronal sections revealed that *EfnA5* was highly expressed in the developing barrel cortex, especially in deep layers, at P7. *Rorβ* marks the barrel field (dashed outline). **(B)** *EfnA5* is expressed at very low levels in a different part of the developing cortex, the anterior motor region, where *Rorβ* expression is also very low. **(C)** Model addressing the role of EfnA5-EphA6 interaction in the reduction of contralateral barrel field innervation downstream of *Mef2c* LOF in S1-L2/3 CPNs. Under this model, we hypothesized that depletion of EfnA5 from the contralateral barrel field should partially restore innervation by *Mef2c^FL/FL^* S1-L2/3 CPNs from the ipsilateral side. **(D)** Double-IUE experimental workflow to (i) knockdown EfnA5 in the contralateral barrel cortex, and (ii) label and knockout *Mef2c* in S1-L2/3 CPNs of the ipsilateral cortex, in the same brain. **(E** and **F)** High-magnification image of the contralateral barrel field from E12.5 contra- *shScrambled* (E) or contra-*shEfnA5* (F) double-IUE brains. GFP+ cell bodies, which express shRNA constructs, are broadly distributed throughout the deep layers (E’’, F’’). tRFP+ axons from *Mef2c^FL/FL^* mutant ipsilateral S1-L2/3 neurons innervated a larger area in the *shEfnA5* cortex (F’, F’’’) than in the *shScram* cortex (E’, E’’’). **(G** and **H)** Coronal sections from double-IUE brains showing *Mef2c^FL/FL^* S1-L2/3 CPNs labeled by tRFP, and expressing *shScrambled* (G, G’) or *shEfnA5* (H, H’) in the contralateral cortex. Mutant neurons displayed low cS1-S2 innervation in the control condition (asterisk, G’), but increased innervation upon *EfnA5* knockdown at the target (arrow, H’). **(I** and **J)** cS1-S2 innervation by *Mef2c^FL/FL^* S1-L2/3 neurons, normalized to ipsilateral cell body intensity, was significantly increased upon *EfnA5* knockdown in the contralateral cortex compared to *shScrambled* expression (I). Ipsilateral E15.5 electroporation efficiency, measured as tRFP signal intensity of ipsilateral L2/3 cell bodies from low-exposure images, was not significantly different between the two groups (J). Data are presented as median +/- IQR; whiskers represent range. n=5 contra-*shScrambled*-Control, 2 litters; and 8 contra-*shEfnA5*, 3 litters. Mann-Whitney U-test: p=0.0016 (I) and 0.5237 (J). ns, not significant; **, p<0.01. Scale bars: 500 μm (A, B, F’’’, H’), 2000 μm (H) See also Figure S7.

To test this prediction, we conducted double-IUE experiments in *Mef2c^FL/FL^* embryos. First, at E12.5 we electroporated shRNA constructs targeting *EfnA5*, or scrambled shRNA controls, into one brain hemisphere with the goal of depleting *EfnA5* across all cortical layers. This was followed at E15.5 by IUE in these same embryos of constructs expressing Cre and a cell-label in the contralateral hemisphere, deleting *Mef2c* in L2/3 CPNs and then observing axon development of these S1-L2/3 CPNs (Figure 7D). We verified with HCR *in situ* hybridization that the E12.5 IUE of *shEfnA5* leads to depletion of *EfnA5* in the electroporated cortex, including in deep layers (Figures S7D-S7G’). We also confirmed that double-IUE appropriately targets deep layers of the E12,5 electroporated side, as well as labeling callosal neurons on the E15.5 electroporated side (Figures 7E-7F’’). Double-IUE in *Mef2c^FL/+^* mice did not result in any differences in *Mef2c^FL/+^* S1 L2/3 CPN axon targeting to *shEfnA5+* contralateral cortex, as compared to *shScram+* contralateral cortex (Figures S7A-S7C). This observation is consistent with our model, in which control S1-L2/3 CPN axons are initially insensitive to EfnA5 repulsion and therefore loss of EfnA5 at the contralateral target does not alter their innervation pattern.

However, we observed that *shEfnA5* expression in contralateral S1 leads to increased innervation of that same contralateral cortical domain by ipsilateral *Mef2c^FL/FL^* S1 L2/3 CPNs, as compared to the innervation of *shScram-*electroporated S1 cortex by *Mef2c^FL/FL^* mutant ipsilateral S1-L2/3 CPNs (Figures 7G-7J). This observation demonstrates that *Mef2c* mutant S1-L2/3 CPN axons are indeed sensitive to repulsion by EfnA5 at contralateral homotopic targets since the relief of repulsion by target-specific *EfnA5* knockdown restores target innervation by *Mef2c* mutant CPN axons.

Together with our observations involving *EphA6* downregulation in *Mef2c^FL/FL^* S1-L2/3 CPNs, our results show that downregulation of *EphA6* by *Mef2c* is necessary to prevent L2/3 CPN axons from being repelled by EfnA5 in the homotopic contralateral target domain.

## DISCUSSION

In this study, we report a novel role for *Mef2c* in mediating the targeting of callosal projections from the somatosensory cortex (S1) to homotopic domains of the contralateral cortex. Postnatal expression of *Mef2c* is enriched in callosal projection neurons (CPNs), and using *in utero* electroporation-based strategies we have developed that allow deletion of *Mef2c* specifically in S1-L2/3 CPNs combined with robust labeling of axonal projections, we reveal strongly impaired projection to contralateral homotopic targets following *Mef2c* LOF.

*MEF2C* LOF variants are strongly linked to neurodevelopmental and neuropsychiatric disorders^38^, and the expression of many axon guidance genes is dependent on Mef2c^34,37^. We establish here, to our knowledge, the first functional link between the dysregulation of a specific axon guidance gene (*EphA6*) downstream of *Mef2c* deletion and a corresponding deficit in interhemispheric connectivity by demonstrating that: (1) overexpression of *EphA6* in S1-L2/3 CPNs reduces their contralateral innervation, phenocopying *Mef2c* LOF; and (2) downregulation of EphA6 function in *Mef2c-*mutant S1-L2/3 CPNs significantly restores contralateral innervation. We also confirm that *EphrinA5 (EfnA5)*, which encodes the repulsive ligand for EphA6, is highly expressed in the somatosensory domain and we observe that downregulation of *EfnA5* in the target cortex significantly restores contralateral innervation by *Mef2c* mutant S1-L2/3 CPNs. Therefore, by downregulating *EphA6* levels Mef2c enables appropriate interpretation of *EfnA5* areal expression to enable homotopic innervation by S1-L2/3 CPNs.

Our results further underscore the significance of repulsive guidance cues in shaping topographic innervation maps^79^. Area specific expression of the repulsive cue *EfnA5*, coupled with complementary expression of EphA receptors, establishes distinct domains that permit contralateral homotopic innervation while repelling projections from heterotopic domains. Interestingly, patterned expression the *EfnA5* ligand also regulates area-specific cortical innervation by thalamic axons^56,58^. The establishment of ‘go/no-go’ zones by a single cue, combined with differential responsiveness imparted by variation in receptor levels between neuronal subtypes, illustrates how a limited number of cell-surface molecules can orchestrate assembly of complex and highly specific circuit wiring patterns.

In *Mef2c* conditional mutants, contralateral CPN targeting is prominently reduced as early as P8, a time when wild type CPN axons first innervate their homotopic cortical domains. The loss of contralateral innervation persists into adulthood and is accompanied by a loss of connectivity in target regions. While other transcription factors like Satb2 and NeuroD2/6 have been shown to regulate callosal projection elaboration, transcription factors imparting specific effects on homotopic targeting after midline crossing remain to be discovered. Therefore, transcriptional programs downstream of Mef2c offer a first glimpse into the molecular logic that dictates patterns of homotopic targeting of callosal projections.

The initial targeting deficit observed in *Mef2c*-mutant CPNs appears distinct from defects arising due to impairment of developmental neuronal activity L2/3 CPNs^9,80–83^. In these cases, initial contralateral innervation is minimally altered at P8, and loss of innervation is apparent only in the second postnatal week. These observations distinguish *Mef2c*-dependent initial homotopic targeting of callosal axons from activity-dependent refinement and selective axon process elaboration in the second postnatal week. On the other hand, sensory-evoked activity-dependent strengthening of L4-to-L2/3 local, ipsilateral, synaptic connectivity requires *Mef2c* in both post-synaptic L2/3 neurons and pre-synaptic L4 neurons^36,37^. Whether the functions of Mef2c in regulating long-range CPN connectivity and in activity-dependent strengthening of local ipsilateral synapses influence each other remains an open question.

Axon-axon interactions have been observed to affect topographic sorting of axons within the corpus callosum^17,19^, and matched cell-surface adhesion among axons of homotopic ipsilateral and contralateral cortical regions has been suggested to influence axon targeting^18^. Our results indicate that target-derived cues also regulate callosal projection innervation, and they identify EfnA5 as one of the first such examples. Indeed, the observation that midline crossing and extension of axons within the white matter is minimally affected in *Mef2c* mutants, and that only the innervation of the appropriate region of the cortical plate is altered, support the idea that EfnA5 exerts its effect at the target rather than through axon-axon interactions along the tract. S1 L2/3 CPNs are sensitive to EfnA5 repulsion only upon *Mef2c* deletion, and *Mef2c LOF* does not impact the targeting of WT S1-L2/3 CPN axons, which express low levels of *EphA6*. What function might ephrinA5 perform during the normal development of homotopic callosal axon projections? EphA-receptor expression is high in more medial cortical domains, including cingulate and retrosplenial cortices^60^, and callosal axons from these regions avoid contralateral motor and somatosensory domains. It is likely that high levels of *EfnA5* in the somatosensory and motor cortical domains function to repel heterotopic projections from these more medial regions.

The EphB2 receptor also functions as a target derived cue can regulate S1-L2/3 CPN contralateral innervation^53,84^. NMDA-dependent clustering of EphB2 on cortical neuron dendrites restricts CPN axon targeting to the S1-S2 border and prevents excessive innervation of the barrel field^53,84^. This is thought to be mediated by callosal axon ephrin B reverse signaling, but it remains unclear whether it involves initial targeting similar to what is mediated by EfnA5, or later functions involving activity-dependent refinement of contralateral projections in the second postnatal week.

Our manipulation of EfnA5-EphA6 signaling downstream of *Mef2c* significantly, but only partially, restored contralateral CPN innervation, suggesting that additional mechanisms promoting homotopic S1 cortex innervation operate downstream of *Mef2c*. Further investigation of additional candidate cell-surface molecules whose expression is regulated in *Mef2c* mutant cortical neurons may identify additional downstream regulators, including those that directly promote callosal axon innervation of S1. It will also be important to investigate intracellular signaling and cytoskeletal regulators that regulate callosal axon projection, including candidates such as Kif2c, which has been shown to promote axon branching downstream of Mef2c^30^.

In addition to contralaterally projecting axons, S1-L2/3 harbors two additional major populations of long-range intracortical projection neurons that project to ipsilateral M1 and to ipsilateral S2, respectively^1,85,86^. Deletion of *Mef2c* also caused target innervation defects in these projections. However, unlike the complete loss of innervation of the contralateral domain, *Mef2c* mutant L2/3 neurons show laminar-specific deficits in the innervation of only the superficial, but not deep, layers of ipsilateral long-range targets. Though our manipulations of EphrinA-EphA signaling downstream of *Mef2c* did not rescue ipsilateral long-range targeting deficits, the remaining cell-surface molecules whose expression is deregulated in the *Mef2c*-mutant cortex^34,37^ offer an opportunity to discover novel regulators of long-range intracortical axon targeting.

*Mef2c-*mutant S1 L2/3 CPNs also displayed ectopic subcortical innervation, including robust projections to the ipsilateral basolateral amygdala (BLA). Given the crucial role of the amygdala in social and emotional processing^87^, and the association of neurodevelopmental disorders with amygdala function defects^88^, it will be interesting to investigate the behavioral consequences of S1 CPN ectopic projections to the BLA.

In addition to defining a novel role for Mef2c in postnatal callosal projection targeting, we shed new light on the role of Mef2c in cortical neuron differentiation during embryogenesis. We localized the embryonic function of *Mef2c* to postnatal cortical neurons, not neural progenitors, and we also observe a novel requirement for post-mitotic *Mef2c* in the development of Rorβ*+* cortical L4 neurons. Reduced *Rorβ* mRNA expression in the embryonic cortex of *Nex-Cre; Mef2c^FL/FL^* mutants foreshadows the absence of any Rorβ+ neurons in the S1 cortex at postnatal stages. We did not detect elevated cell death upon pan-cortical deletion of *Mef2c* at embryonic or early postnatal stages, suggesting that L4 neurons, rather than being lost to cell death, are mis-specified into a different class of neuron lacking expression of many cortical laminar subtype markers.

Rorβ+ L4 neurons are a characteristic feature of primary sensory cortical areal organization, distinguishing S1 from more frontal/motor areas^89^. Pilot assessments of cortical areal marker expression in *Nex-Cre; Mef2c^FL/FL^* brains revealed that, concomitant with the Rorβ+ L4 neuron loss, there is a downregulation of S1-areal markers and upregulation of frontal/motor markers in *Nex-Cre; Mef2c^FL/FL^* mutants (data not shown). Future investigations will shed light on how laminar-subtype specification and areal identity acquisition are coordinated by Mef2c during post-mitotic cortical neuron development.

Taken together, our work highlights the multifunctional roles served by Mef2c during cortical development: in the embryonic development of cortical laminar organization and in postnatal homotopic targeting of callosal projections. Importantly, we identify a novel role for EfnA5-EphA signaling in orchestrating callosal projection targeting downstream of Mef2c. These results offer a first glimpse into the molecular logic underlying homotopic organization of callosal projections. *MEF2C* loss-of-function is strongly associated with neurodevelopmental and neuropsychiatric disorders^38,39^, as is aberrant intracortical connectivity that includes disrupted callosal projection homotopy^14–16^. Our characterization of Mef2c as a regulator of intracortical projection targeting suggests that the molecular programs regulated by MEF2C will provide crucial insights into the etiology of aberrant interhemispheric connectivity in neurodevelopmental and neuropsychiatric disorders, with the potential for offering therapeutic intervention targets. Moreover, since *MEF2C* is highly expressed in other brain regions during development, including in regions of the cerebellum, hippocampus and lateral amygdala^38,60^, this work may help address roles played by Mef2c more widely in the regulation of axon projection targeting across multiple brain regions.

### Limitations of this study

Though we show that downregulation of *EphA* receptor expression by Mef2c is necessary for homotopic innervation by S1-L2/3 CPNs, it remains to be determined whether *EphA6* transcription is directly repressed by Mef2c or if a different Mef2c target downregulates *EphA6* expression. Mef2c can function as both a transcriptional repressor and activator; repressor function is crucial in establishing excitatory/inhibitory synaptic balance during cortical development^34^, and activator function modulates activity-dependent strengthening of L4->L2/3 connectivity^37^. Functional assessment of potential Mef2c-binding sequences in close to the *EphA6* TSS will address whether it is a direct target of Mef2c.

## MATERIALS AND METHODS

### EXPERIMENTAL MODEL DETAILS

#### Mice

Experiments were carried out in strict accordance with the recommendations in the *Guide for the Care and Use of Laboratory Animals* of the National Institutes of Health. The animal protocol was approved by the Animal Care and Use Committee of The Johns Hopkins University School of Medicine (Protocol #M023M68). Mice of both sexes were used in all experiments. Noon of the day after the plug was designated as E0.5, and the date of birth as P0. Mice used in all experiments were group housed and maintained in a 14/10-hour (h) light/dark (LD) cycle and had access to food and water *ad libitum*.

*Mef2c-floxed* mice in the C57BlL6/J background (*Mef2c^tm1Jjs^*/J, Jax Strain #025556), kindly shared by John Schwarz^62^, were crossed to WT CD1 mice (Charles River Laboratory) and maintained on a mixed background as heterozygotes and homozygotes. *Nex-Cre (Neurod6-Cre)^tg/tg^* mice in the C57BlL6/J background, a gift from Klaus-Armin Nave^61^, were crossed to mixed-background *Mef2c^Fl/Fl^* mice to generate *Mef2c^Fl/+^; Nex-Cre^Tg/+^* stock. Timed pregnant CD1 females were obtained from Charles River Laboratory (Strain code 022) for WT IUE experiments.

### METHODS

#### Oligonucleotide Sequences

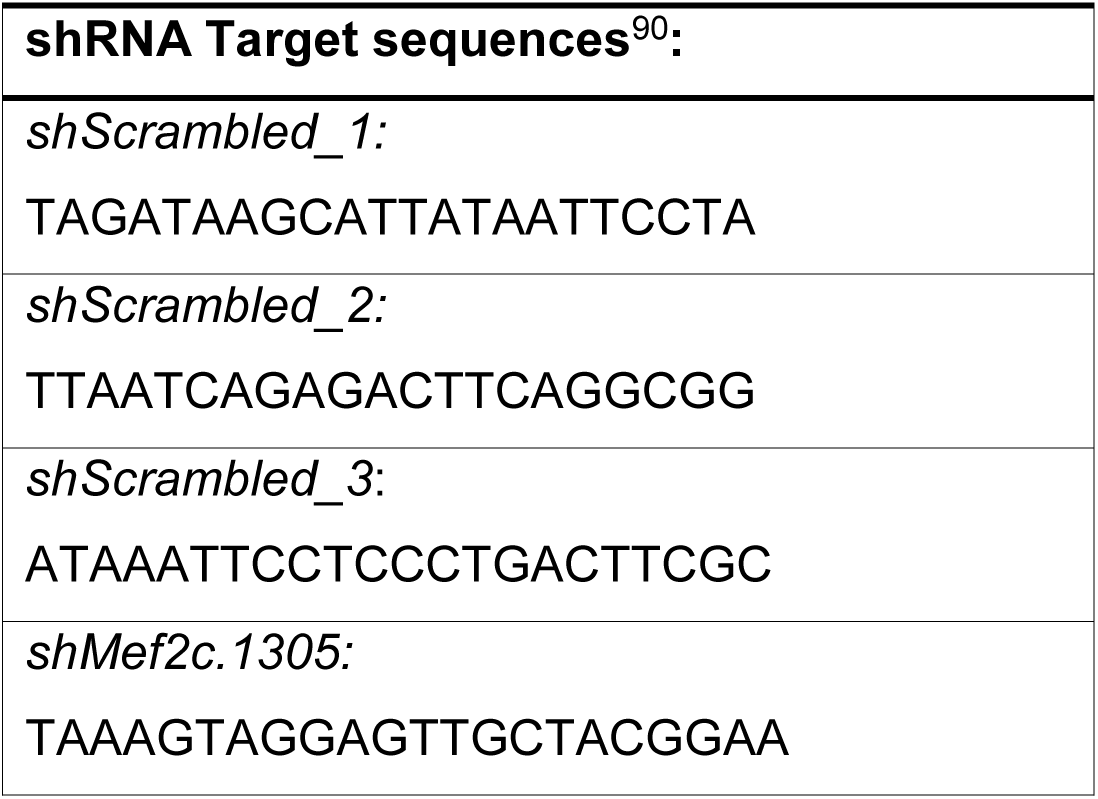
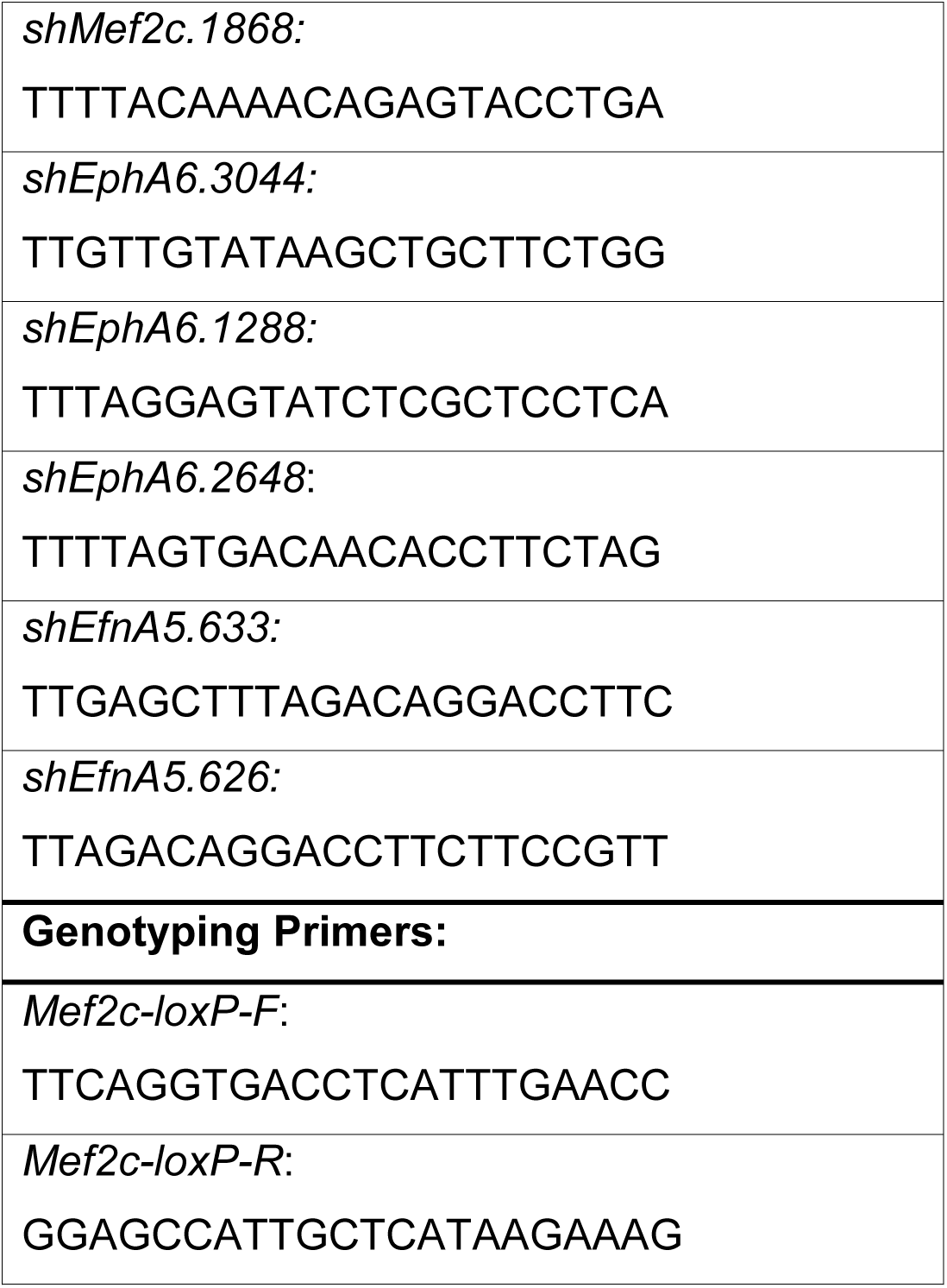

#### Plasmids

*pDCX-Cre-ires-mCherry,* modified from *pDCX-Cre-ires-GFP* ^66^, was kind gift from Soraia Barao and Ulrich Mueller, and *pCβA-Flex*^69^ was a gift from Ulrich Mueller. *pTRE-Cre* (Addgene #69136) and *pCAG-Lox-Stop-Lox-turboRFP-ires-tTA-WPRE* (*pCAG-LSL-tRFP-irestTA,* Addgene #69138)^67^, as well as *pCAG-Lox-Stop-Lox-EGFP-WPRE* (*pCAG-LSL-GFP-ires-tTA,* Addgene #85006) and *pCAG-FRT-Stop-FRT-turboRFP-ires-tTA-WPRE* (*pCAG-FSF-tRFP-ires-tTA,* Addgene #85038) were gifts from Takuji Iwasato. *pCAFNF-EGFP* (Addgene #13722)^91^ was a gift from Constance Cepko, *pTRE- DIO-FlpO* (Addgene #118027)^74^ from Minmin Luo, *pCAG-Flex-mWGA-mCherry-WPRE* (*pCAG-Flex-mWmC*)^70^ from Xin Duan, and *pCAG-FlpO* (Addgene #125576)^92^ from Takeshi Imai.

To generate the *pCAG-FSF-EphA6-HA* overexpression construct, the C-terminal HA-tagged *EphA6* cDNA ORF (Sino Biological MG50630-CY) was excised as a KpnI-NotI fragment and cloned in to the corresponding sites of *pCAFNF-*EGFP, replacing the GFP cassette.

To generate the *pCAG-FSF-EphA6ΔIntraCellularDomain(ICD)-GFP*, a PCR fragment including the N-terminal extracellular region and transmembrane domain and only the first 4 amino acids of the intracellular domain was amplified from the full-length *EphA6* cDNA ORF (Sino Biological MG50630-CY), fused with an in-frame *EGFP* sequence in an intermediate cloning vector (Randal Hand and Alex Kolodkin, unpublished), and then cloned into the XhoI-NotI sites of *pCAG-FSF-ires-GFP*^72^, replacing the *ires-GFP* cassette.

To generate the *pCAG-FSF-GFP-shmiRE* cloning vector, the *miR30* backbone was excised as an NotI-MscI fragment from *pPRIME-dsRed-shScram*^93^ and cloned into the corresponding sites of *pCAG-FSF-GFP*^72^, downstream of the GFP cassette. The *FSF* cassette was excised to generate the Flp-independent pCAG-GFP-*shmiRE* vector. *shRNA* target sequences for the genes of choice were selected from a previous study^90^, and 97-bp oligomeric DNA corresponding to *miR30E*-*shRNA* sequence, as specified in Fellman *et al* 2013^90^, were synthesized. Oligomers were PCR amplified with *XhoI-site*-containing forward and *EcoRI-site-*containing reverse primers to generate inserts that were then cloned into the XhoI-EcoRI sites of the appropriate vector (*pCAG-FSF-GFP- shmiRE* for *shEphA6*, and *pCAG-GFP-shmirE* for *shMef2c* and *shEfnA5*). The target sequences of all *shRNAs* used in this study are listed in the Oligonucleotide Sequences table above.

All plasmids generated in this study were validated by sequencing, and the NucleoBond Xtra Midi EF kit (Takara Bio 740422.50) was used for endotoxin-free preparation of all plasmids used for IUE.

#### In Utero Electroporation

*In utero* electroporation to target S1 L2/3 or deep layer neuronal progenitors were performed, on E15.5 or E12.5 timed-pregnant mice, as previously described^72,73,78^. Briefly, time-pregnant mice were anesthetized with 2% isoflurane and placed on a heating pad. The abdomen was shaved and cleaned, 100uL of the local anesthetic Bupivacaine-hydrochloride (2.5 mg/mL; Sigma B5274-5G) was applied, and a 1.5-2 cm laparotomy was made. Pups were extracted and rinsed with sterile warmed PBS. Pups were injected in one lateral ventricle using glass needles pulled on a vertical pipette puller (Narishige PC-10), and electroporated with three 40 V pulses (E15.5) or five 36 V pulses (E12.5) of 50 ms duration delivered at 1 Hz, administered using a BTX ECM 830 square pulse electroporator (Harvard Apparatus) via 5mm gold-plated paddle electrodes (BTX 450170, Fischer Scientific) at E15.5 or CUY650P3 3-mm round tweezer electrodes (NepaGene) at E12.5. DNA constructs were mixed in PBS with Fast Green FCF dye (Sigma F7258-25G) to visualize injections. 0.75 µL of Buprenorphine (1 mg/mL; ZooPharm LLC) was administered for post-operative pain control.

The following plasmid combinations were used in this study:

For S1 L2/3 CPN-specific *Mef2c* conditional knockout, *Mef2c^Fl/+^* E15.5 timed pregnant females (crossed with *Mef2c^Fl/+^* males) were injected with *pDcx-Cre-ires-mCherry* (0.6 µg/µL) and either *pCAG-LSL-tRFP-ires-tTA* (1.0 µg/µL) or *pCAG-LSL-GFP-ires-tTA* (1.5 µg/µL). *mCherry* is no longer expressed from the *DCX*-promoter at all postnatal stages analyzed in this study, and only the other, Cre-dependent, fluorophore is detected.

For dual labeling, *Mef2c^FL/+^* E15.5 timed pregnant females (crossed with *Mef2c^FL/+^* males) were injected with *pTRE-Cre* (0.125 µg/µL), *pβA-Flex* (1.5 µg/µL) and *pCAG-LSL-GFP-ires-tTA* (1.5 µg/µL).

For S1 L2/3 CPN-specific *Mef2c* knockdown in WT CD1 E15.5 timed pregnant females, half the embryos were injected with *pCAG-mCherry* (1.5 µg/µL) and a pool of 2 *pCAG- GFP-shMef2c* plasmids, each expressing a different *Mef2c* targeting *shRNA* (0.75 µg/µL each); the rest of the embryos were injected in the opposite hemisphere with *pCAG-mCherry* (1.5 µg/µL) and a pool of 2 *pCAG-GFP-shScram* plasmids, each expressing a different scrambled-control *shRNA* (0.75 µg/µL each).

For WGA-anterograde tracing, *Mef2c^FL/+^* E15.5 timed pregnant females (crossed with *Mef2c^FL/+^* males) were injected with *pDcx-Cre-ires-mCherry* (0.6 µg/µL), *pCAG-Flex-mWmC (*1.5 µg/µL) and *pCAG-LSL-GFP-ires-tTA* (1.5 µg/µL).

For *EphA6* overexpression, *pDcx-Cre-ires-mCherry* (0.6 µg/µL) *pTRE-DIO-FlpO* (0.6 µg/µL) and *pCAG-FSF-tRFP-ires-tTA* (1 µg/µL) were mixed with either *pCAG-FSF-EphA6-HA (*1.5 µg/µL) or *pCAFNF-GFP* (1.5 µg/µL). In WT CD1 E15.5 timed pregnant females, half the embryos were injected, in the same hemisphere, with *EphA6-HA* mix, while the other half were injected, in the other hemisphere, with *GFP* mix.

For *EphA6ΔICD* expression, *pDcx-Cre-ires-mCherry* (0.6 µg/µL) *pTRE-DIO-FlpO* (0.6 µg/µL) and *pCAG-FSF-tRFP-ires-tTA* (1 µg/µL) were mixed with either *pCAG-FSF-EphA6ΔICD-GFP (*1.0 µg/µL) or *pCAFNF-GFP* (1.0 µg/µL). In *Mef2c^FL/+^* E15.5 timed pregnant females (crossed with *Mef2c^FL/FL^* males), half the embryos were injected, in the same hemisphere, with the mix containing *EphA6ΔICD-GFP*, while the other half were injected in the other hemisphere with control mix that contained *pCAFNF-GFP* instead.

For *EphA6* knockdown, *pDcx-Cre-ires-mCherry* (0.6 µg/µL) *pTRE-DIO-FlpO* (0.6 µg/µL) and *pCAG-FSF-tRFP-ires-tTA* (1 µg/µL) were mixed with either a pool of three *pCAG-FSF-GFP-shEphA6* plasmids, each expressing a different *EphA6* targeting *shRNA* (0.6 µg/µL each), or with a pool of three *pCAG-FSF-GFP-shScram* plasmids, each expressing a different scrambled-control *shRNA* (0.6 µg/µL each). In *Mef2c^FL/+^* E15.5 timed pregnant females (crossed with *Mef2c^FL/FL^* males), half the embryos were injected in the same hemisphere with *shEphA6* mix, while the other half were injected in the other hemisphere with *shScram* mix.

For shRNA validation, either the *pCAG-FSF-GFP-shEphA6* pool (3 x 0.6 µg/µL) or the *pCAG-FSF-GFP-shScram* pool (3 x 0.6 µg/µL) was mixed with *pCAG-FlpO* (0.6 µg/µL) and *pCAG-FSF-EphA6-HA (*1.5 µg/µL). In WT CD1 E15.5 timed pregnant females, half the embryos were injected in the same hemisphere with *shEphA6* mix, while the other half were injected in the other hemisphere with *shScram* mix.

For *EfnA5* knockdown by double IUE, all embryos in *Mef2c^FL/FL^* E15.5 timed pregnant females (crossed with *Mef2c^FL/+^* males) were first injected in the same hemisphere with a pool of two *pCAG-GFP-shEfnA5* plasmids (0.75 µg/µL each) for the Contra-*shEfnA5* condition, or, with a pool of two *pCAG-GFP-shScram* plasmids (0.75 µg/µL each) for the Contra-*shScram* condition. Three days later, at E15.5, the same timed pregnant mothers were subjected to a second IUE. At this stage all embryos were injected in the hemisphere opposite to E12.5 injection with a mix of *pDcx-Cre-ires-mCherry* (0.6 µg/µL) *pTRE-DIO-FlpO* (0.6 µg/µL) and *pCAG-FSF-tRFP-ires-tTA* (1 µg/µL). The same double IUE experiments were also performed with *Mef2c^FL/+^* timed pregnant females crossed with *Mef2c^+/+^* males.

To validate efficiency of *EfnA5* knockdown, CD1 WT E12.5 timed pregnant females were injected in a single hemisphere with the pool of two *pCAG-GFP-shEfnA5* plasmids (0.75 µg/µL each).

#### Immunohistochemistry in thick brain sections to visualize axon projections

Mice younger than P7 were deeply anesthetized on ice, and mice P7 or older were deeply anesthetized with CO2. They were then transcardially perfused with ice-cold phosphate-buffered saline (PBS), followed by 4% paraformaldehyde (Electron Microscopy Sciences #15711) in PHEM buffer (27 mm PIPES (Amresco; 0169-250G), 25 mm HEPES (Sigma; H3375-500G), 5 mm EGTA (Amresco; 0732-1006), 0.47 mm MgCl2 (Sigma; M8266-100G), pH 6.9) with 10% sucrose and 0.1% Triton X-100 (4% PFA/PHEM). Brains were dissected and evaluated for IUE efficiency under a fluorescence dissection microscope. Brains with very low signal intensity or very high IUE signals outside the S1 region were discarded at this stage. Brains with good, S1-specific IUE signal were post-fixed in 4% PFA/PHEM overnight (O/N) at 4^0^C. Fixed brains were washed in PBS 3 times, 1 hour each, and stored in sealed containers at 4^0^C for until sectioning. 250-μm coronal sections were prepared using a vibratome (Leica VT1000s). Sections were then washed 3x in PBS, and treated with permeabilization solution (0.1% Triton X-100 and 3% Bovine Serum Albumin) in PBS for 1 hour at RT, and incubated overnight at 4°C in blocking solution (permeabilization solution + 5% Normal Goat Serum (NGS, Jackson ImmunoReseach 005-000-121) with primary antibodies. The following primary antibodies were used at the indicated concentration: rabbit anti-tagRFP (Thermo Fisher MA5-32668), 1:1000; chicken anti-GFP (Aves GFP-1020), 1:1000; rabbit anti-dsRed (Takara Bio 632496); 1:1000; rat anti-HA (Sigma 11867423001), 1:500; rabbit anti-Mef2c (Cell Signaling 5030S), 1:1000. Sections were then washed 3x in PBS and incubated overnight at 4^0^C in PBS + 0.1% Triton X-100 with secondary antibodies and DAPI (Fisher Scientific D1306). Finally, sections were washed 3x in PBS, mounted with Aqua Polymount (Polyscience Inc. 18606-20) on Superfrost Plus^TM^ (Fisher Scientific 12-55-15) slides, and then stored at 4^0^C until imaging. The following cross-adsorbed secondary antibodies were used at a 1:1000 dilution: goat anti-rabbit Alexa Fluor 555 (Invitrogen A21429); goat anti-chicken Alexa Fluor 488 (Invitrogen A32931); goat anti-rabbit Alexa Fluor 647 (Invitrogen A21245); and goat anti-rat Alexa Fluor 488 (Invitrogen A48262).

#### Immunohistochemistry and In Situ Hybridization Chain Reaction (HCR) in Thin Coronal Brain Sections

P7-8 mice E16.5 timed pregnant females were deeply anesthetized with CO2, and P0 mice with ice. They were then transcardially perfused with ice-cold, RNase-free phosphate-buffered saline (PBS), followed by RNase-free 4% PFA in PBS. Brains were then isolated from embryos and pups, and post-fixed overnight in RNase-free 4% PFA-PBS overnight at 4^0^C. Brains were then washed in PBS 3 times, 1 hour each, and cryoprotected overnight in RNase-free PBS + 30% Sucrose at 4^0^C. The next day, tissue was frozen over dry ice in NEG-50 (Thermo Fisher 22-046-511) blocks. Fixed-frozen brains were cryosectioned at thickness of 16µm on a Leica CM3050 cryostat. Sections spanning the entire somatosensory cortex were collected on Superfrost Plus^TM^ (Fisher Scientific 12-55-15) slides. Fixed-frozen tissue blocks and cryosection slides were stored at -80^0^C until use.

HCR probe hybridization and amplification were carried out according to manufacturer’s protocol (molecularinstruments.com) for fixed-frozen cryosections. Hybridization was performed overnight with HCR probes designed and synthesized by Molecular Instruments against the following *Mus musculus* mRNA sequences: *Mef2c (*compatible with HCR-hairpins B2)*, Rorβ (B1), Pax6 (B1), EphA6 (B5), EphA7(B4)* and *EfnA5 (B2)*. 20 pairs of probes, each at a concentration of 4nM, were used per mRNA. This was followed by overnight room-temperature amplification with HCR-hairpins coupled to AlexaFluor 546 or 647, compatible with the appropriate initiators, at a concentration of 60nM. Slides were then washed with 5X SSCTw (Saline-Sodium Citrate Buffer + 0.1% Tween-20) before proceeding to immunohistochemistry (IHC). If only HCR was required, slides were counterstained with DAPI and mounted with Aquapolymount after 5X SSCTw washes.

For IHC, sections were first blocked with 10% NGS in PBS+0.1%Triton for 1 hour at RT followed by overnight 4^0^C staining with primary antibodies in the same blocking solution. The following primary antibodies were used on thin sections: Chicken anti-GFP (Aves GFP-1020), 1:500; Rabbit anti-Brn2 (Cell Signaling 12137S), 1:1000; Mouse anti-Rorβ (Perseus Proteomics PP-N7927-00), 1:250; Rat anti-Ctip2 (Abcam ab18465), 1:1000; Rabbit anti-Cux1 (Proteintech 11733-1-AP), 1:500; Rabbit anti-Cux2 (Proteintech 24902-1-AP), 1:500; Rabbit anti-Cleaved Caspase 3 (Cell Signaling 9661), 1:500; Rabbit anti-Bhlhb5 (Abcam ab204791), 1:1000 and Guinea Pig anti-Vglut2 (Sigma Aldric ab2251-I), 1:500. Sections were then washed 3x in PBS at RT and incubated for 2hr at RT in PBS + 0.1% Triton X-100 with secondary antibodies and DAPI (Fisher Scientific D1306). Finally, sections were washed 3x in PBS and mounted with Aquapolymount (Polyscience Inc. 18606-20) and stored at 4^0^C until imaging. The following cross-adsorbed secondary antibodies were used at a 1:500 dilution: goat anti-rabbit Alexa Fluor 647 (Invitrogen A21245), goat anti-rabbit Alexa Fluor 488 (Invitrogen A11034), goat anti-rabbit Alexa Fluor 555 (Invitrogen A21429), goat anti-chicken Alexa Fluor 488 (Invitrogen A32931), goat anti-rat Alexa Fluor 488 (Invitrogen A48262), goat anti-rat Alexa Fluor 647 (Invitrogen A21247) and goat anti-guinea pig Alex Fluor 647 (Invitrogen A21450).

#### Confocal Imaging of Brain Sections

All imaging was performed on a Zeiss LSM700 confocal microscope controlled by Zen 2012 SP5 software (Zeiss).

Tiled images of entire thick coronal sections were acquired with a 10X objective (NA 0.3) at 1.25 µm x 1.25 µm pixel size in xy and z-intervals of 20 µm, spanning the entire tissue depth. Two separate images were acquired of each section, one at high laser intensity and gain to visualize axon projections, and a second at lower laser intensity and gain to obtain images where ipsilateral L2/3 cell body fluorescence was not saturated. Laser power and gain settings for the low and high intensity conditions were kept constant for all brains that were quantified and compared as part of any single experiment.

Tiled images of the cS1-S2 border region in EGFP-tdTomato dual-labeled brains were obtained with a 20X (NA 0.8) objective at 0.31 µm x 0.31 µm pixel size in xy and z-interval of 1.5 µm, spanning the entire tissue depth. A single-tile image of the corpus callosum at the midline, also at 0.31 µm x 0.31 µm in xy and z-interval of 1.5 µm, spanning the entire tissue depth was obtained for normalization. The same laser settings were applied to both types of images and across all brains.

Tiled images of the cS1-S2 border region in mWGA-mCherry and GFP labeled brains were also obtained with the 20X (NA 0.8) objective at 0.31 µm x 0.31 µm pixel size in xy and z-interval of 1.5 µm, up to a depth of 12 µm. The same laser settings were applied to all brains. A second, low-magnification, low intensity tiled image of the ipsilateral cortex was obtained with the 10X objective. The same setting was applied to low-mag low-intensity images of all brains. The fluorescence intensity of ipsilateral L2/3 GFP cell bodies from these images was used as measure of IUE efficiency.

Zoomed images of L2/3 cell bodies were acquired for *Eph* receptor HCR and co-expression analysis of L2/3 and S1 markers in electroporated neurons with a 40X oil-immersion objective (NA 1.3) at 0.08 µm x 0.08 µm pixel size in xy and z-interval of 1 µm, spanning the entire tissue depth. Four coronal sections, separated from each other by 125 µm and spanning the S1 barrel region were imaged for each brain. Care was taken to ensure matching of anterior-posterior sections among different brains based on DAPI-staining of landmarks. Three images were acquired of each section, at approximately matched locations progression from the barrel field medial start to the S1-S2 border. Laser power and gain settings were kept constant for all brains that were quantified and compared as part of an experiment.

For analysis of cortical laminar organization, tiled images, spanning the Pia-to-WM axis of the barrel cortex in height and ∼600 µm (P7) or ∼300 µm (E16.5) in width, were obtained with a 20X (NA 0.8) objective at 0.31 µm x 0.31 µm pixel size in xy and z-interval of 1.5 µm, spanning the entire tissue depth. Three to four coronal sections separated from each other by 125 µm (P7) or 80 µm (E16.5) and spanning the S1 barrel region were imaged for each brain. Care was taken to ensure matching of anterior-posterior sections among different brains based on DAPI-staining of landmarks. Laser power and gain settings were kept constant for all brains that were quantified and compared as part of an experiment. Images used for display were cropped to a reduced width.

Tiled images of entire brain sections were obtained with a 10X (NA 0.3) objective at 1.25 µm x 1.25 µm pixel size in xy and z-interval of 3 µm.

### QUANTIFICATION AND STATISTICAL ANALYSIS

All image analyses were performed with FIJI/ImageJ^94^ unless stated otherwise.

#### Quantification of contralateral innervation and midline crossing

To quantify raw axon innervation of the cS1/S2 border, the integrated density in a manually drawn region of interest (ROI) was first obtained from high-intensity coronal section images of electroporated brains. The horizontal extent of the ROI covered he last two visible barrels in S1, and an equivalent distance in S2. Vertically, the ROI covered all cortical layers up to the pia, but excluded sub-cortical white matter. DAPI staining was used to determine the vertical and horizontal boundaries. At P8, the ROI was drawn to cover the entire contralateral barrel field and a small portion of S2 close to the S1-S2 border. The integrated fluorescence intensity in this ROI was treated as the raw innervation.

To account for variability due to electroporation, we next obtained the integrated density of an ROI spanning all labeled cell-bodies in ipsilateral L2/3 from low intensity images of the same coronal section. The cS1/S2 integrated density measure was then divided by the ipsilateral L2/3 cell body intensity to obtain the normalized cS1/S2 innervation value.

Using DAPI staining of anatomical landmarks, we ensured that matched coronal sections were analyzed across all brains in a single experiment. Analysts were blinded to genotype and/or IUE-based perturbation condition.

Quantification of midline crossing was also performed from maximum intensity projection images of coronal sections. Integrated fluorescence intensity was calculated in three manually drawn ROIs in the pre-crossing, midline, and post-crossing portions of the corpus callosum and treated as raw values. These raw values were then then divided by the ipsilateral L2/3 cell body intensity from low-intensity images for normalization.

#### Innervation profile analysis

Innervation profiles along the Pia-to-WM axis of cS1-S2 and the medial-to-lateral axis of the contralateral barrel field were first generated using the plot profile tool of Fiji/ImageJ by analysts who were blind to the genotype. A vertical line spanning the Pi-to-WM axis was extended into a rectangle of width 500 µm to cover the cS1-S2 border domain, and the intensity profile along the vertical axis was generated. For intensity profiles along the horizontal axis of the barrel field, a horizontal segmented line spanning the barrel field and cS1-S2 was extended to form a polygon of width of 750 µm.

Raw intensity profiles were exported as .csv files and processed with a custom Python script that normalized the total distance from 0 to 1 (WM to Pia in the vertical profiles, and medial end to lateral end in the horizontal profiles). The script was then used to calculate interval sums of intensity in 10 equally spaced intervals across the axis and to plot the mean and standard deviation at each interval for each condition analyzed in the experiment. Statistical comparisons were also performed using the script: first testing the values at each interval for homogeneity of variance (Levene’s test), and then followed by an unpaired T-Test or a Mann-Whitney U-test, as appropriate. The p-values obtained for each interval were adjusted for multiple comparisons with a Bonferroni correction and adjusted p-values <0.05 were considered statistically significant.

#### Quantification of EGFP:tdTomato intensity ratio in dual-labeled brains

A custom-written Fiji/ImageJ macro was used to sum the total intensity, across all z-planes, of individual channels of composite .czi z-stack images. This macro was employed on composite images of cS1-S2 or the corpus callosum midline of brains from dual labeling experiments to calculate the ratio of GFP to tdTomato channel intensity. During imaging, DAPI-staining of anatomical landmarks was used to select matching coronal sections across different brains for analysis.

#### WGA cell counting

DAPI staining was used to locate the cS1-S2 border region from tiled 20x z-stack images of the contralateral cortex. L5 and L2/3 were next delineated in this region and also based on DAPI staining. The total number of nuclei with perisomatic-WGA signal were counted across individual z-planes in 300 µm-wide rectangular regions of cS1-S2 L5 and cS1-S2 L2/3 and then summed to obtain the raw total contralateral WGA count.

Next, electroporation efficiency was calculated as the ipsilateral L2/3 GFP signal from low-intensity 10x images of ipsilateral cortex of the same coronal section for which contralateral WGA was quantified. Raw WGA count was then divided by ipsilateral L2/3 GFP intensity to obtain the normalized cC1-S2 WGA cell count. Analysts were blind to genotype, and matching coronal sections were analyzed across all brains in the experiment.

#### HCR/ IHC analysis of electroporated L2/3 neurons

*EphA* receptor mRNA HCR signal was quantified from high-magnification z-stack images of electroporated L2/3 neurons by an analyst blind to genotype. 3-5 neurons were chosen at random from each image for analysis. An ROI corresponding to the neuron-cell body (marked by electroporated GFP) was delineated. This was done in the z-plane where the cell-body area was largest for the chosen neuron *EphA6* HCR signal intensity as well as the number of puncta within the ROI were recorded.

To quantify Brn2, Cux1, Bhlhb5 and CC3 co-expression, electroporated L2/3 neurons and their appropriate z-plane were also chosen as described for HCR quantification. They were then classified as positive (1) or negative (0) for the marker being assessed, based on the IHC signal. Then, the total number of positive neurons was calculated for each condition.

#### HCR/ IHC analysis of cortical laminar markers

Maximum projections of tiled images of P7 images of S1 cortex were first cropped to width of 150 µm, while maintaining the entire height from Pia-to-WM. The total number of Rorβ+, Ctip2+ and Cux2+Rorβ- neurons were counted within this cropped region, and the Pia-to-WM distance (cortical plate thickness) was also measured in the DAPI channel. Counts and cortical thickness were obtained from images taken from four coronal sections separated from each other by 125 µm and spanning the anterior-posterior extent of the barrel cortex, and then averaged for each brain. Analysts who performed the above operations were blinded to animal genotypes. This was then followed by litter normalization, where the average value for all control (*Mef2c^FL/+^; Nex- Cre*) animals in a litter was normalized to 1.

Analysis of E16.5 S1 cortex laminar organization was performed similarly with a few modifications. The cropped area used for counting was 100µm wide, and Brn2+ and Ctip2+ cell numbers and total DAPI nuclei were counted in this area. In addition, total *RorB* HCR signal intensity within the cortical plate was measured. Images from three coronal section, separated from each other by 80 µm, were quantified and averaged for each brain. Values were then normalized to the control (*Mef2c^FL/+^; Nex-Cre*) mean of the litter.

#### Statistical analysis

All graphs were generated and statistical tests were performed using Prism version 10 (GraphPad) unless otherwise specified. Individual animals were considered as biological replicates (n), and all experiments involve comparisons to littermate controls except for the double-IUE experiments. The number of animals used (n), statistical measures represented in graphs, statistical tests performed, and p-values are all reported in the legends. The threshold for statistical significance was defined as p < 0.05.

### RESOURCE AVILABILITY

#### Lead contact

Further information and requests for resources and reagents should be directed to and will be fulfilled by the lead contact, Alex L. Kolodkin.

#### Materials availability

Plasmids generated in this study will be shared by the lead contact upon request

#### Data and code availability

- Custom Python code and Fiji/ImageJ Macros will be made available publicly on GitHub.
- Any additional information required to reanalyze the data reported in this paper is available from the lead contact upon request.

## ACKNOWLEDGMENTS

We are grateful to Dr. John Schwarz and Dr. Klaus-Armin Nave for kindly sharing mouse strains, and Drs. Randal Hand, Soraia Barao, Ulrich Mueller, Xin Duan, Takuji Iwasato, Constance Cepko, Takeshi Imai and Minmin Luo for their kind gifts of plasmids. We thank Dr. Ulrich Mueller, Dr. Solange Brown, Dr. Soraia Barao, Yijun Xu, Dr. Fengquan Zhou and all members of the Kolodkin laboratory for insightful discussion and suggestions. We are especially grateful to Nicole Kropkowski and Dr. Victoria Neckles for excellent technical assistance and laboratory management. This work was supported by the Distinguished Graduate Student Fellowship from the Johns Hopkins Kavli Neuroscience Discovery Institute to S.S, Johns Hopkins Neuroscience Scholars Program Fellowship to L.G.C., and R03-TR004616 and Johns Hopkins School of Medicine Institutional Funds to A.L.K.

## AUTHOR CONTRIBUTIONS

Conceptualization, S.S., and A.L.K.; methodology, S.S., L.G-C., and A.L.K.; investigation, S.S., L.G-C., N.D., and N.S.; writing—original draft, S.S., L.G-C., and A.L.K.; writing— review & editing, S.S., L.G-C., J.Z., A.L.K.; funding acquisition, S.S., L.G-C., and A.L.K.; resources, S.S, J.Z., and X.O.J.; supervision, A.L.K.

## SUPPLEMENTAL INFORMATION

**Document S1. Figures S1–S7**

