## Supplemental Data 1 for "Mef2c Controls Postnatal Callosal Axon Targeting by Regulating Sensitivity to Ephrin Repulsion"

### Figure S1, related to Figure 1

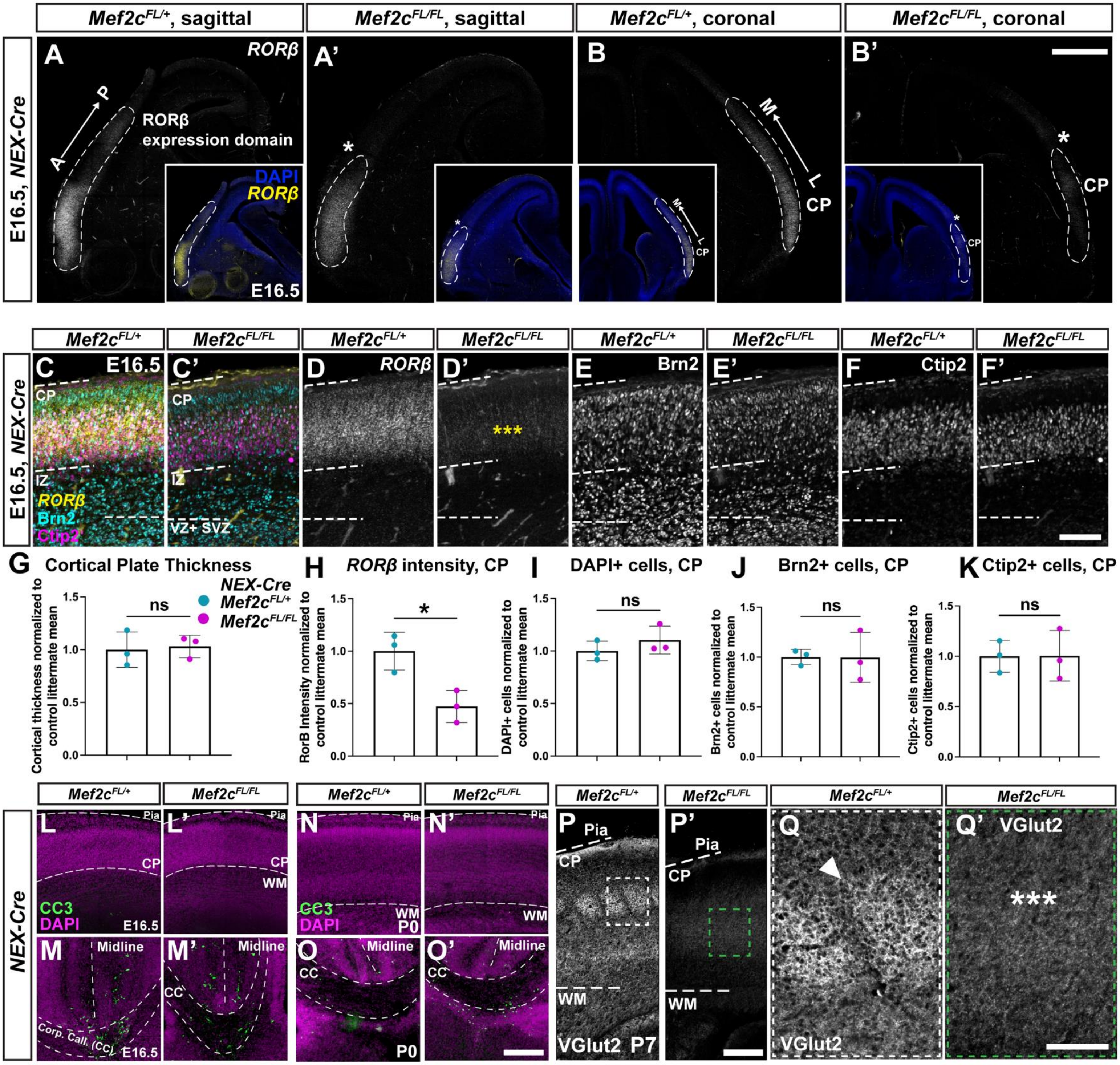

**Figure S2, related to Figure 1**

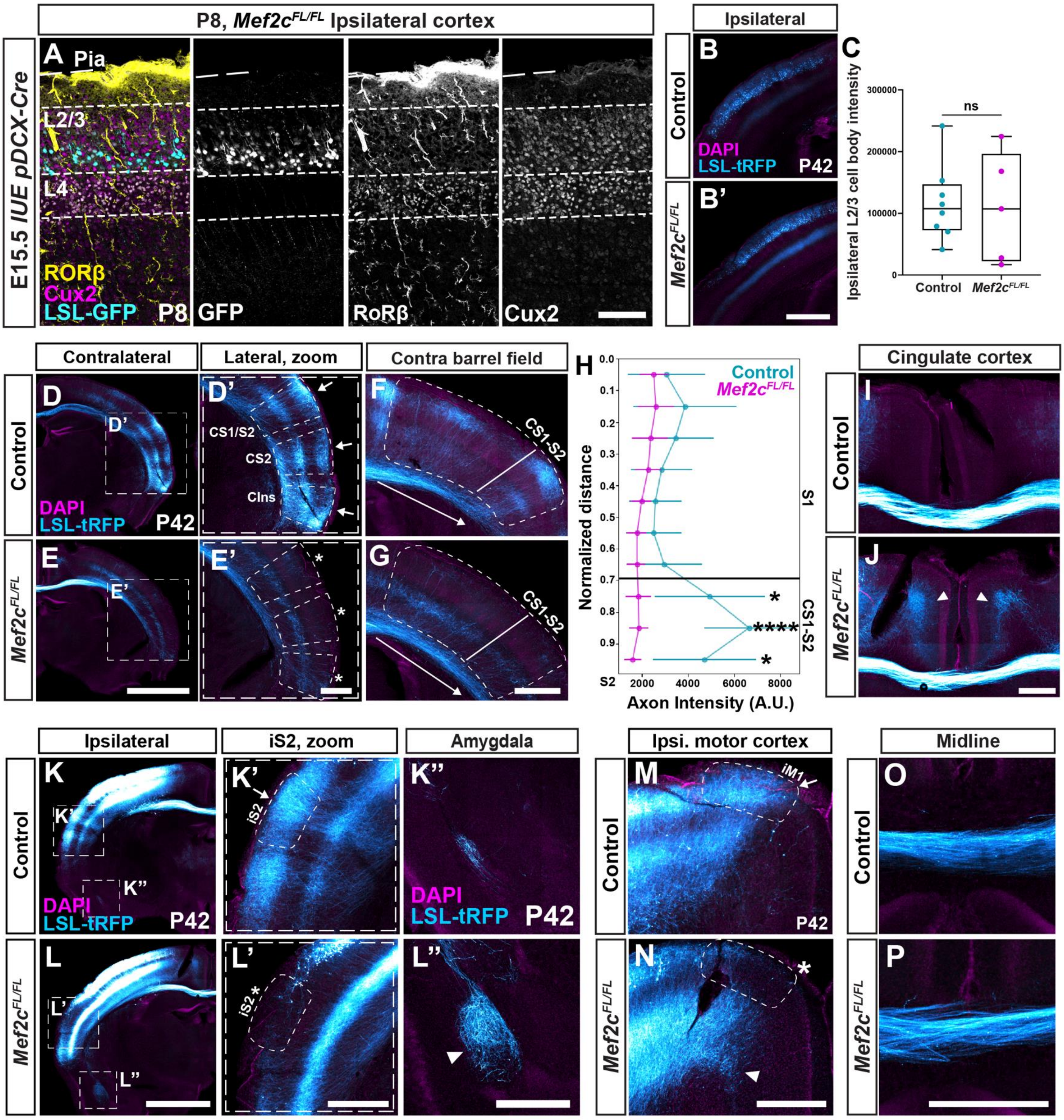

**Figure S3, related to Figure 2**

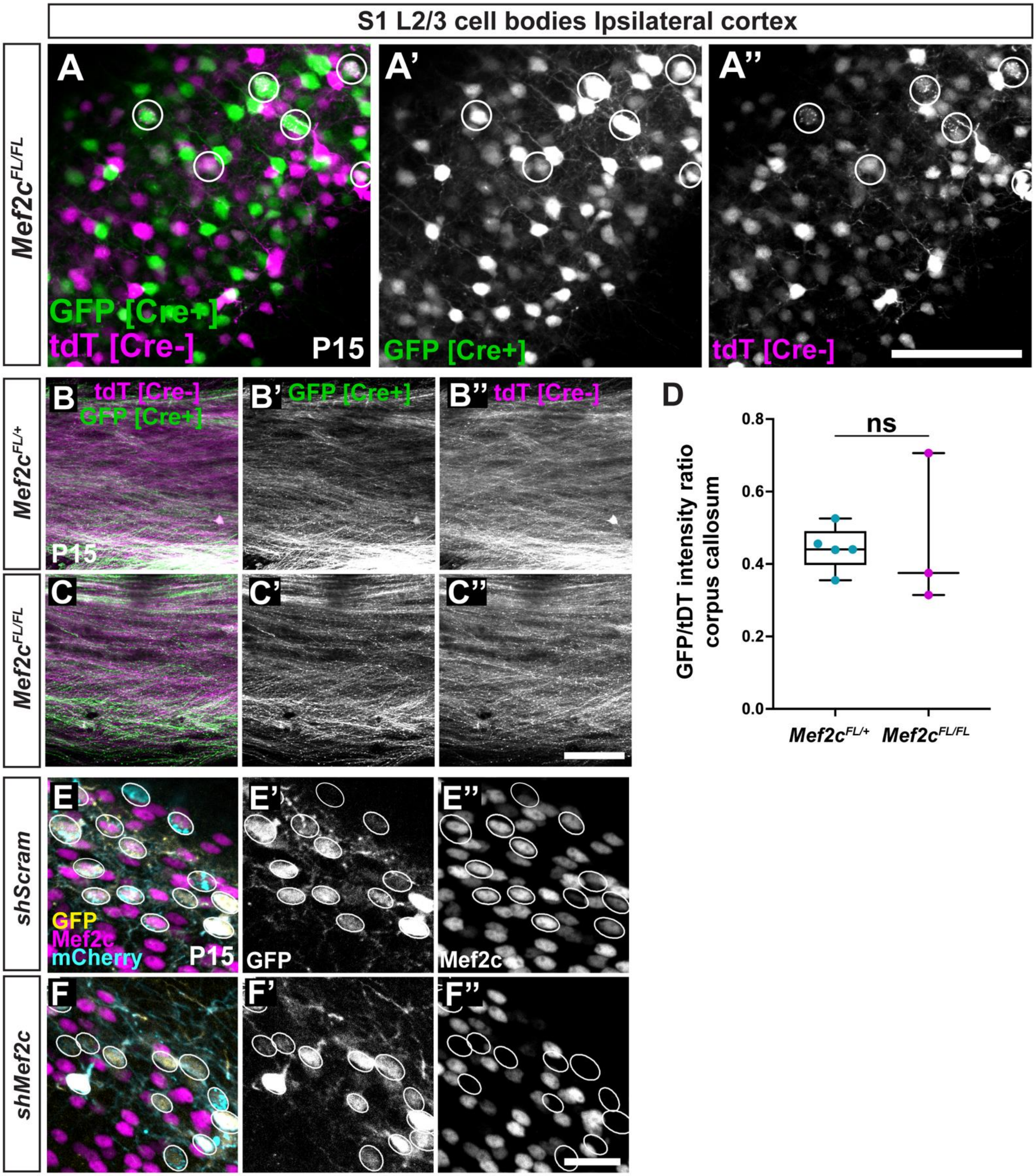

### Figure S4, related to Figure 4

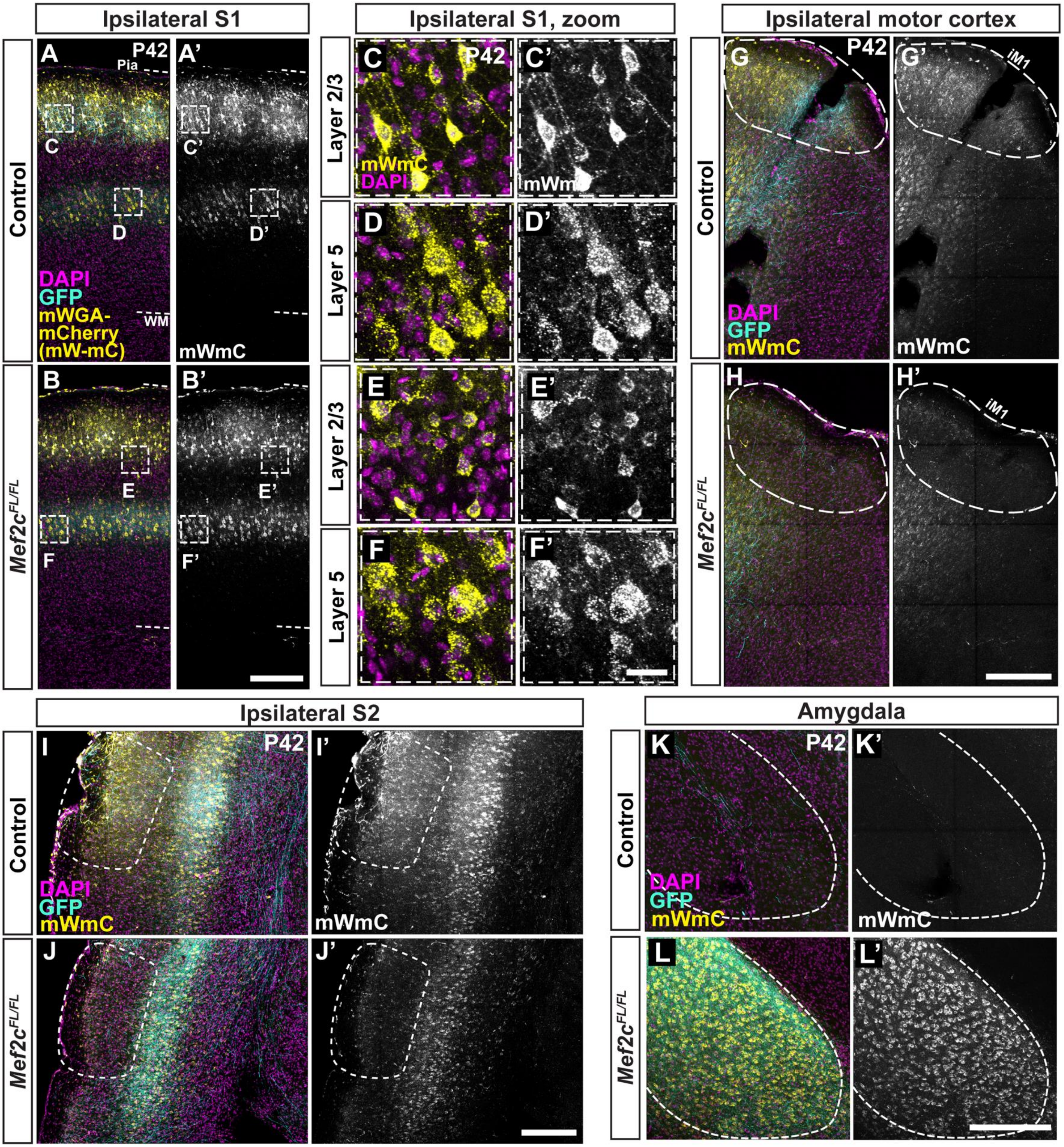

### Figure S5, related to Figure 5

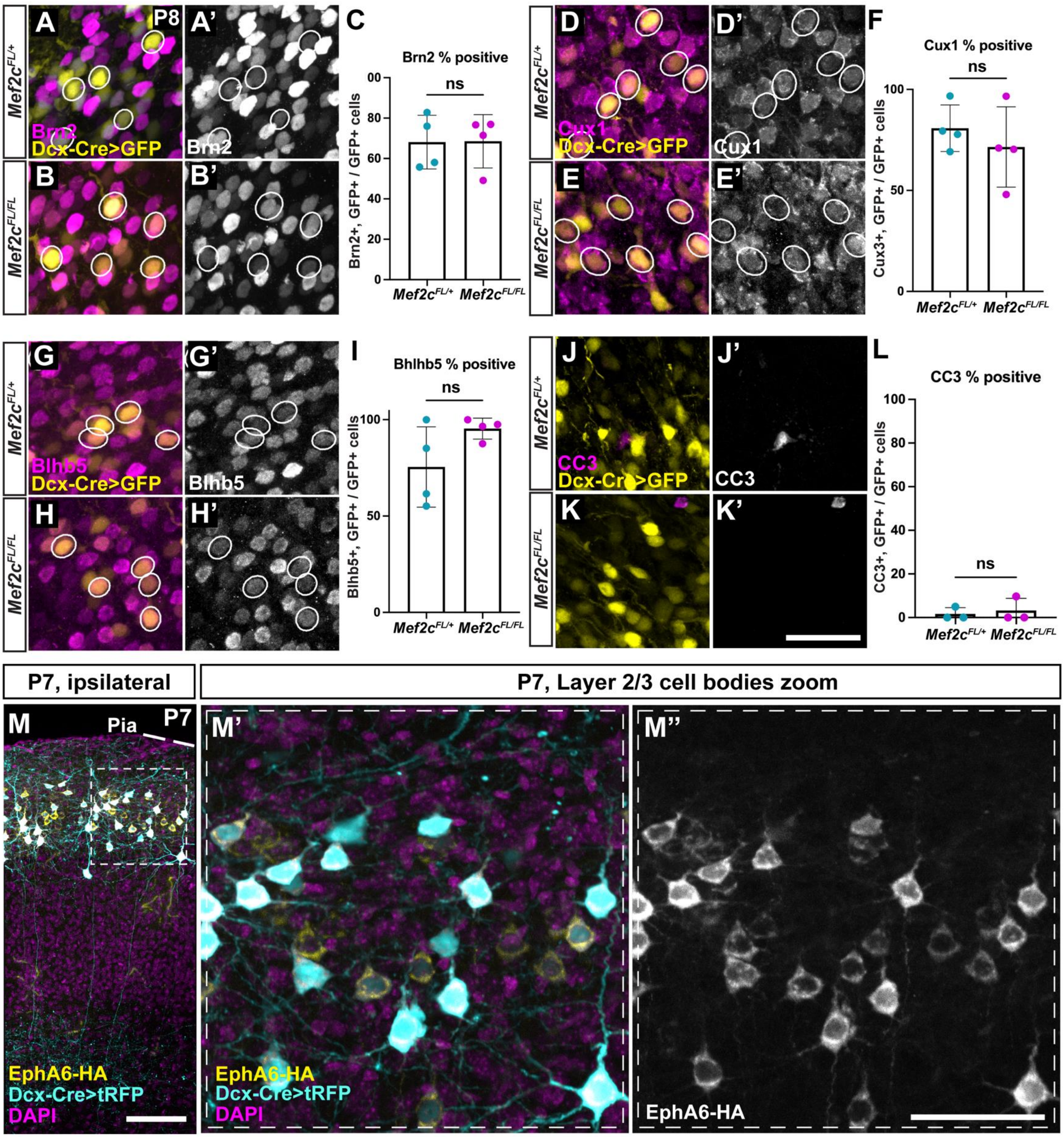

#### Figure S6, related to Figure 6

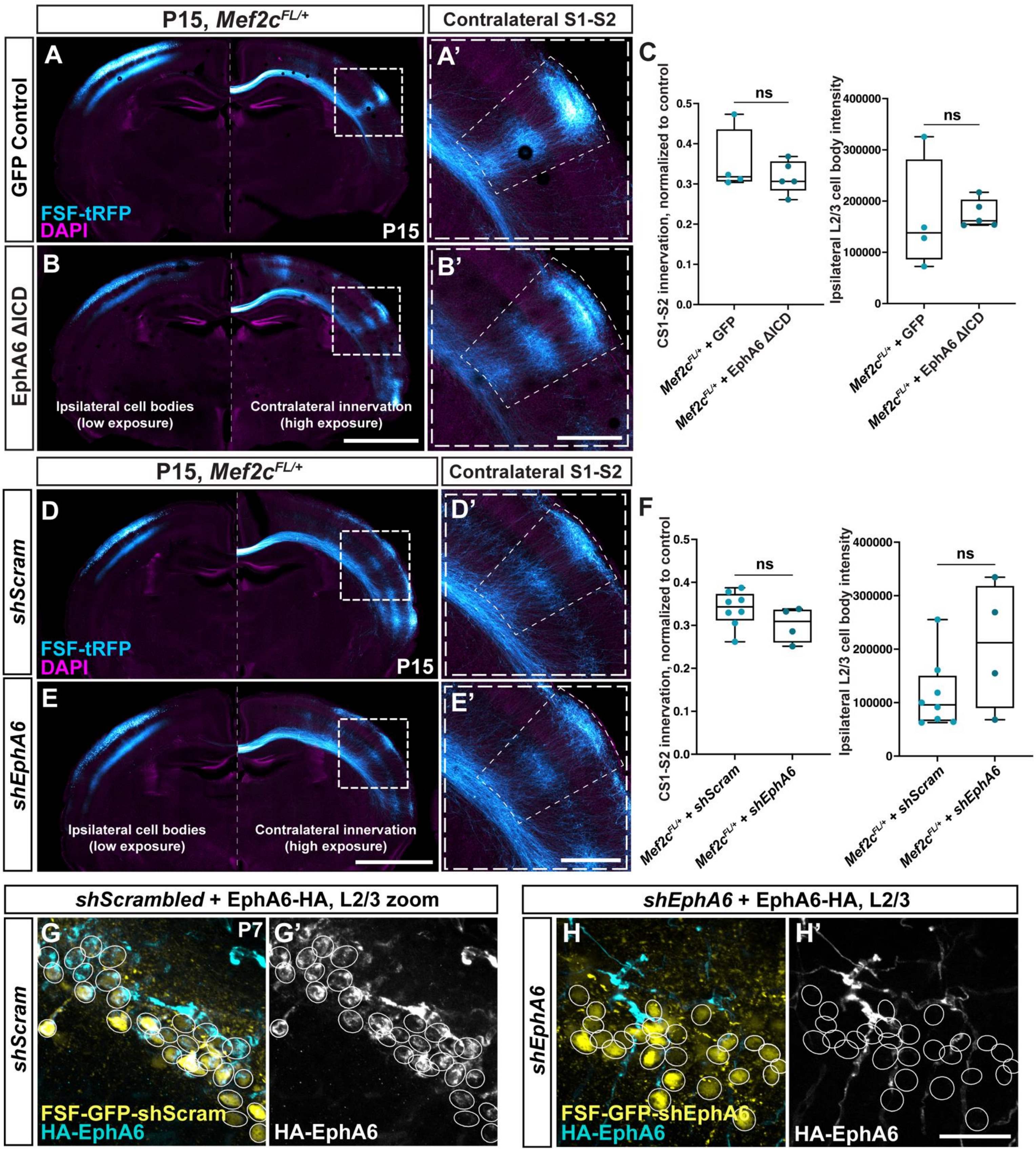

### Figure S7, related to Figure 7

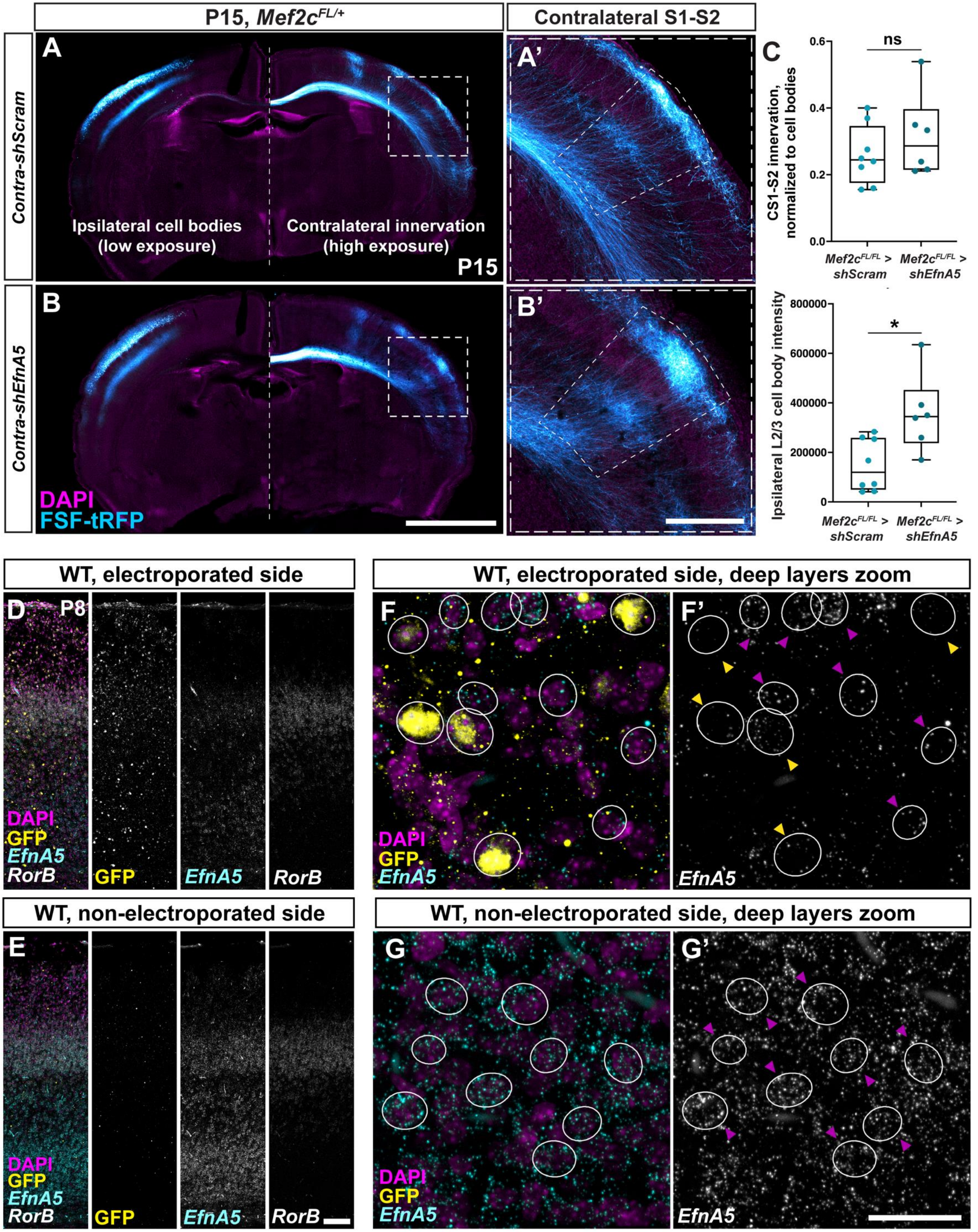

**Figure S1 (Related to Figure 1): *Rorβ* expression in the embryonic cortical plate is reduced upon pan-cortical, post-mitotic deletion of *Mef2c***

**(A** and **B)** The *Rorβ* mRNA expression domain in the cortical plate (dashed lines) is greatly reduced upon pan-cortical *Mef2c* deletion driven by *Nex-Cre*, as seen in both sagittal (*$Mef2c^{FL/+}$* Control in A and *$Mef2c^{FL/FL}$* in A') and coronal (B and B') sections at E16.5. Insets show composite images, with nuclear DAPI staining.

**(C-F):** Coronal sections through S1 of E16.5 *Nex-Cre; $Mef2c^{FL/+}$* (C, D, E, F) and *Nex-Cre; $Mef2c^{FL/FL}$* mutant (C', D', E', F') littermates, labeled with laminar subtype markers. *Rorβ* mRNA expression is reduced in the *Nex-Cre; $Mef2c^{FL/FL}$* mutant cortical plate at E16.5 (yellow asterixis in D'), while Brn2 (E, E') and Ctip2 (F, F') are unaffected.

**(G-K)** In *Nex-Cre; $Mef2c^{FL/FL}$* mice CP thickness (G), the number of DAPI nuclei (I), Brn2 nuclei (J) and Ctip2 nuclei (K) in the CP were all unaffected, while *Rorβ* mRNA intensity (H) was reduced compared to *Nex-Cre; $Mef2c^{FL/+}$*littermates at E16.5. Graphs represent mean +/- s.d, n=3 animals/condition, 3 sections per animal. Values were normalized to control littermate means. Unpaired two-tailed Student's T-test, p=0.7979 (G) 0.0183 (H), 0.3292 (I), 0.9855 (J), 0.9820 (K).

**(L-O)** Cleaved-Caspase 3 (CC3) staining is not detectable in *Nex-Cre; $Mef2c^{FL/+}$* control or *Nex-Cre; $Mef2c^{FL/FL}$* mutant cortical plates at E16.5 (L, L') and P0 (N, N'), suggesting the absence of elevated cell death in mutant cortices. CC3+ cells were detectable at the midline and corpus callosum in both genotypes at both ages (M, M' and O, O').

**(P** and **Q)** Staining for Vglut2 reveals thalamocortical afferent segregation into barrels in the P7 *Nex-Cre; $Mef2c^{FL/+}$* control cortex (P, arrowhead in Q) but not in the *Nex-Cre; $Mef2c^{FL/FL}$* mutant cortex (P', asterisks in Q').

Scale bars: 500 µm (B'), 200 µm (F', P', O'), 100 µm (Q')

**Figure S2 (Related to Figure 1): Additional characterization of *Mef2c*-mutant S1-L2/3 CPN targeting deficits**

**(A)** *Mef2c$^{FL/FL}$* brains electroporated with Cre at E15.5 show normal migration of Cre+ neurons to L2/3, as well as normal expression of upper layer markers Cux2 (L2/3 + L4) and Rorβ (L4), at P8.

**(B)** Low-exposure images of ipsilateral electroporated region of Control (B) and *Mef2c$^{FL/FL}$* (B') brains, Similar number and distribution of tRFP+ cell bodies in ipsilateral L2/3 at P42 are apparent in both conditions.

**(C)** Quantification of tRFP intensity in ipsilateral L2/3, a measure of IUE efficiency, from low exposure images, reveals similar distributions for Control and *Mef2c$^{FL/FL}$* brains at P42. Data are presented as median +/- IQR; whiskers represent range. n=8 control and 5 *Mef2c$^{Fl/Fl}$* mice, 3 litters. Mann-Whitney U-test: p=0.8329.

(**D** and **E**) Additional contralateral innervation deficits of *Mef2c$^{FL/FL}$* S1-L2/3 neurons at P42. Insets are magnified regions lateral to the contralateral barrel field (BF), revealing reduced innervation in all three domains (cS1-S2, cS2 and contralateral Insular cortex=cIns) by *Mef2c$^{FL/FL}$* neurons (asterisks, E') compared to controls (arrows, D').

**(F** and **G)** Innervation in the middle of the contralateral BF was affected to a lesser degree than at the S1-S2 boundary in *Mef2c$^{FL/FL}$* mutants (F vs G). Dashed boxes denote the BF; solid lines, start of cS1-S2; arrows, axis along which intensity profiles in F were calculated.

**(H)** Comparison of innervation fluorescence intensity profiles from Control and *Mef2c$^{Fl/FL}$* littermates along the dorsomedial to ventrolateral (DM-to-VL) axis of the contralateral BF, showing no significant differences in any regions other than the S1-S2 boundary at the ventrolateral end at P42. Data are presented as mean +/- s.d of innervation intensity in bins that correspond to 1/10$^{th}$ of the normalized BF extent. n=8 control and 5 *Mef2c$^{FL/FL}$* mice, 3 independent litters. Unpaired T-test with Bonferroni correction for multiple comparisons, p=0.0405, 0.0006 and 0.0290 at 0.75, 0.85 and 0.95 normalized BF DM-to-VL distance, respectively.

**(I** and **J)** *Mef2c*$^{FL/FL}$ S1-L2/3 CPNs exhibit ectopic axonal projections in deep layers of the cingulate cortex, both ipsilaterally and contralaterally (arrowheads, J), which are absent in Controls (I) at P42.

**(K** and **L)** *Mef2c*$^{FL/FL}$ S1-L2/3 CPNs show laminar specific innervation deficits in ipsilateral S2 (iS2) and ectopic innervation of the amygdala. Control neurons innervate both deep and superficial layers of iS2 (arrow, K'), while *Mef2c*$^{FL/FL}$ neurons fail to innervate superficial layers (asterisk, L'). *Mef2c*$^{FL/FL}$ S1-L2/3 CPNs also show ectopic innervation of the basolateral amygdala (arrowhead, L'').

**(M** and **N)** *Mef2c*$^{FL/FL}$ S1-L2/3 CPNs also show laminar specific innervation deficits in the ipsilateral motor region (iM1). Control neurons innervate both deep and superficial layers of iM1, while *Mef2c*$^{FL/FL}$ neurons fail to innervate superficial layers (asterisk, N). Mutant neurons also mis-project to regions more medial to iM1 (arrowhead, N).

**(O** and **P)** Midline crossing is unaffected upon *Mef2c* LOF in S1 L2/3 CPNs, as seen from images of the corpus callosum of Control (O) and *Mef2c*$^{FL/FL}$ (P) electroporated brains at P42.

ns, not significant; *, $p<0.05$; ****, $p<0.001$.

Scale bars: 200 µm (A) 500 µm (B', E', G, J, L', L'', N, P), 2000 µm (E, L)

**Figure S3 (Related to Figure 2): Dual-labeling and *Mef2c* knockdown validation**

**(A)** Images of ipsilateral S1-L2/3 neurons upon IUE of dual-labeling constructs (see Figure 2A) in a *Mef2c*$^{FL/Fl}$ animal, indicating mostly non-overlapping GFP (Cre+) (A') and tdTomato (Cre-) (A'') cell bodies. The few instances of dual-labeled cell bodies are circled.

**(B** and **C)** Images of the corpus callosum upon IUE of dual labeling constructs. GFP (Cre+) intensities were comparable between control (B, B') and *Mef2c*$^{Fl/Fl}$ (C, C') brains, as were tdTomato (Cre-) intensities (B, B'' and C, C'').

**(D)** Quantification of Cre+ (GFP): Cre-(tdTomato) axonal intensity ratio in the corpus callosum. No significant differences in the Cre+: Cre- ratio were apparent between *Mef2c*$^{Fl/Fl}$ and *Mef2c*$^{Fl/+}$ littermates. Data are presented as median +/- IQR; whiskers represent range. n=5 *Mef2c*$^{Fl/+}$ and 3 *Mef2c*$^{Fl/Fl}$ mice from one litter. Mann-Whitney U-test, p=0.7857

**(E** and **F)** Validation of *Mef2c* knockdown: WT brains were electroporated with either *pCAG-GFP-shScram* plasmids (E-E'') or *pCAG-GFP-shMef2c* plasmids (F-F'') at E15.5 and immunostained for Mef2c at P15. Mef2c expression was detectable in GFP+ S1-L2/3 neurons of the shScrambled-control group (E', E'''), but absent in GFP+ neurons of the *shMef2c* group (F', F''), indicating efficient Mef2c knockdown. Both groups were also electroporated with the *pCAG-mCherry* cell label.

ns, not significant

Scale bars: 100 µm (A'', C''), 25 µm (F'').

**Figure S4 (Related to Figure 4): Characterization of *Mef2c-cKO* S1-L2/3 CPN synaptic connectivity deficits at additional targets with WGA anterograde tracing**

**(A-F)** Cre-dependent WGA anterograde tracing reveals normal local ipsilateral synaptic connectivity of *Mef2c*$^{FL/FL}$ L2/3-CPNs neurons (B, E and F), similar to controls (A, C and D). Electroporated L2/3 neurons in ipsilateral S1 cortex (GFP+, mWmC+) of Control (A) and *Mef2c*$^{FL/FL}$ (B) both show grossly similar number and pattern of synaptically connected local targets in L2/3 and L5 (GFP-, mWmC+). Insets show magnified images of synaptically connected cell bodies, showing perisomatic mWmC, in Control L2/3 (C), Control L5 (D), *Mef2c*$^{FL/FL}$ L2/3 (E) *Mef2c*$^{FL/FL}$ L5 (F).

**(G and H)** Compared to Control IUE neurons (G), *Mef2c*$^{FL/FL}$ S1-L2/3 neurons are synaptically connected to fewer neurons in the upper layers of ipsilateral motor cortex (H), as visualized by anterograde mWmC transfer.

**(I** and **J)** Compared to Control IUE neurons (I), *Mef2c*$^{FL/FL}$ S1-L2/3 neurons are synaptically connected to fewer neurons in the upper layers of ipsilateral secondary somatosensory cortex (S2) (J), as visualized by anterograde mWmC transfer.

**(K** and **L)** The basolateral amygdala (dashed outline), which is ectopically innervated by *Mef2c*$^{FL/FL}$ S1-L2/3 axons, also displays a large number of mWmC+ cell-bodies that are synaptically connected to *Mef2c*$^{FL/FL}$ S1-L2/3 neurons (L), whereas no perisomatic mWmC was apparent in the amygdala of Control brains (K).

Scale bars: 250 µm (B’, H’, J’, L’), 25 µm (F’)

**Figure S5 (Related to Figure 5): Assessment of L2/3 subtype markers, an S1 areal marker, and cell death in *Mef2c* mutant neurons, and validation of EphA6 overexpression by IUE**

**(A** and **B)** *pDcx-Cre>LSL-GFP* electroporated cell bodies from *Mef2c*$^{FL/+}$ control (A and A') and *Mef2c*$^{FL/FL}$ (B and B') brains, immunostained for the superficial layer (L2/3 +L4) marker Brn2, at P8. Sections from the same mice as in Figure 4B-D, subjected to the same IUE workflow, were used.

**(C)** Quantification of the percentage of GFP electroporated L2/3 neurons that are also positive for the marker Brn2 (Brn2+, GFP+/ GFP+) reveals no differences between *Mef2c*$^{FL/+}$ controls and *Mef2c*$^{FL/FL}$ mutant neurons. Data are represented as mean +/- S.D per mouse. n=171 neurons from N=4 *Mef2c*$^{FL/+}$ mice and 154 neurons from N=4 *Mef2c*$^{FL/FL}$ mice. $p>0.9999$, Mann-Whitney test.

**(D** and **E)** *pDcx-Cre>LSL-GFP* electroporated cell bodies from *Mef2c*$^{FL/+}$ control (D and D') and *Mef2c*$^{FL/FL}$ (E and E') brains, immunostained for the superficial layer (L2/3 +L4) marker Cux1, at P8. Sections from the same mice as in Figure 4B-D, subjected to the same IUE workflow, were used.

**(F)** Quantification of the percentage of GFP electroporated L2/3 neurons that are also positive for the marker Cux1 (Cux1+, GFP+/ GFP+) show no differences between *Mef2c*$^{FL/+}$ controls and *Mef2c*$^{FL/FL}$ mutant neurons. Data are represented as mean +/- S.D per mouse. n= 118 neurons from N=4 *Mef2c*$^{FL/+}$ mice and 113 neurons from N=4 *Mef2c*$^{FL/FL}$ mice. $p=0.2286$, Mann-Whitney test.

**(G** and **H)** *pDcx-Cre>LSL-GFP* electroporated cell bodies from *Mef2c*$^{FL/+}$ control (G and G') and *Mef2c*$^{FL/FL}$ (H and H') brains, immunostained for the S1 areal marker Bhlhb5, at P8. Sections from the same mice as in Figure 4B-D, subjected to the same IUE workflow, were used.

**(I)** Quantification of the percentage of GFP electroporated L2/3 neurons that are also positive for the marker Bhlhb5 (Bhlhb5+, GFP+/ GFP+) shows no differences between *Mef2c*$^{FL/+}$ controls and *Mef2c*$^{FL/FL}$ mutant neurons. Data are represented as mean +/- s.d percentage per mouse. n=105 neurons from N=4 *Mef2c*$^{FL/+}$ mice and 131 neurons from N=4 *Mef2c*$^{FL/FL}$ mice. p=0.4857, Mann-Whitney test.

**(J** and **K)** *pDcx-Cre>LSL-GFP* electroporated cell bodies from *Mef2c*$^{FL/+}$ control (J and J') and *Mef2c*$^{FL/FL}$ (K and K') brains, immunostained for Cleaved Caspase 3 (CC3), an indicator of cell death, at P8. Sections from the same mice as in Figure 4B-D, subjected to the same IUE workflow, were used.

**(L)** Quantification of the percentage of GFP electroporated L2/3 neurons that are also positive for the marker CC3 (CC3+, GFP+/ GFP+) shows no differences between *Mef2c*$^{FL/+}$ controls and *Mef2c*$^{FL/FL}$ mutant neurons. Data are represented as mean +/- s.d percentage per mouse. n=75 neurons from N=3 *Mef2c*$^{Fl/+}$ mice and 81 neurons from N=3 *Mef2c*$^{FL/FL}$ mice. p>0.9999, Mann-Whitney test.

**(M)** Validation of EphA6-HA overexpression by IUE with HA-tag staining. Images of the electroporated cortex (M) and insets of L2/3 (M' and M'') from a mouse subjected to the IUE paradigm in Figure 4G showing the presence of HA-tagged EphA6 in tRFP+ cell bodies.

ns, not significant.

Scale bars: 50 μm (K', M''), 100 μm (M)

**Figure S6 (Related to Figure 6): Assessment of the effects of DN-EphA6 expression and *EphA6* knockdown in *Mef2c*$^{FL/+}$ control neurons by IUE, and validation of *EphA6* knockdown**

**(A** and **B)** *EphA6ΔICD-GFP* expression in *Mef2c*$^{FL/+}$ S1-L2/3 neurons (B, B') does not alter cS1-S2 innervation compared to GFP-expressing *Mef2c*$^{FL/+}$ neurons (A, A'). The same IUE constructs from Figure 5A were introduced into *Mef2c*$^{Fl/+}$ embryos at E15.5 and brains were assessed at P15.

**(C)** Quantification of cS1-S2 innervation, normalized to ipsilateral L2/3 cell body intensity, does not reveal significant differences between *EphA6ΔICD* expressing and GFP- expressing *Mef2c*$^{FL/+}$ S1-L2/3 neurons (Left). Electroporation efficiency, measured as ipsilateral L2/3 cell body intensity from low-exposure images, is comparable between *EphA6ΔICD* IUE and GFP IUE in *Mef2c*$^{FL/+}$ brains (Right). Data are presented as median +/- IQR; whiskers represent range. n=4 GFP-control and 5 EphA6ΔICD brains, 3 litters. Mann-Whitney U-test: p=0.7302 (innervation), p=0.2857 (cell bodies).

**(D** and **E)** *EphA6* knockdown in *Mef2c*$^{FL/+}$ S1-L2/3 neurons (E, E') does not alter cS1-S2 innervation compared to Scrambled control expressing *Mef2c*$^{Fl/+}$ neurons (D, D'). The same IUE constructs from Figure 5G were introduced into *Mef2c*$^{FL/+}$ embryos at E15.5 and brains were assessed at P15.

**(F)** Quantification of cS1-S2 innervation, normalized to ipsilateral cell body intensity, does not reveal significant differences between *shEphA6* and *shScram* expressing *Mef2c*$^{FL/+}$ S1-L2/3 neurons (Left). Electroporation efficiency, measured as ipsilateral L2/3 cell body intensity from low-exposure images, is comparable between *shEphA6* knockdown and *shScram Mef2c*$^{FL/+}$ brains (Right). Data are presented as median +/- IQR; whiskers represent range. n=8 *shScram* and 4 *shEphA6* brains, 4 litters. Mann-Whitney U-test: p=0.2828 (innervation), p=0.2141 (cell bodies).

**(G** and **H)** Validation of *EphA6* knockdown: WT brains were electroporated at E15.5 *pCAG-FlpO* + *pCAG-FSF-EphA6-HA* to overexpress HA-tagged EphA6, combined with either *pCAG-FSF-GFP-shScram* plasmid pool (G, G') or *pCAG-FSF-GFP-shEphA6* plasmid pool (H, H'), and evaluated at P7. GFP+ S1-L2/3 neurons from the *shScram* brain are positive for HA (G, G'), while GFP+ neurons from the *shEphA6* animal are not, indicating efficient knockdown.

ns, not significant.

Scale bars: 2000 μm (B, E), 500 μm (B', E'), 100 μm (H').

**Figure S7 (Related to Figure 7): Assessment of the effects of contralateral *EfnA5* knockdown followed by ipsilateral Cre IUE in *Mef2c*$^{FL/+}$ control neurons (double IUE), and validation of *EfnA5* knockdown in deep cortical layers by shRNA IUE**

**(A** and **B)** *Mef2c*$^{FL/+}$ brains with S1-L2/3 CPNs with tRFP, and expressing *shScrambled* (A and A') or *shEfnA5* (B and B') in the contralateral cortex. cS1-S2 innervation was comparable in both conditions. The same IUE constructs from Figure 6D were introduced into *Mef2c*$^{Fl/+}$ embryos at E12.5 and E15.5 and brains were assessed at P15.

**(C)** cS1-S2 innervation by *Mef2c*$^{FL/+}$ S1-L2/3 neurons, normalized to ipsilateral cell body intensity, was not significantly different upon *EfnA5* knockdown in the contralateral cortex compared to *shScrambled* expression (Top). Ipsilateral L2/3 E15.5 electroporation efficiency, measured as tRFP intensity in ipsilateral L2/3 from low-exposure images, was significantly higher in the contra-*shEfnA5* group (Bottom). Data are presented as median +/- IQR with whiskers representing the range. n=8 contra-*shScram*-control from 2 litters and 6 contra-*shEfnA5* brains from 1 litter. Mann-Whitney U-test: p=0.7546 (innervation), p= 0.0127 (cell bodies)

**(D-G)** Validation of *EfnA5* knockdown in deep cortical layers: *pCAG-GFP-shEfnA5* constructs were electroporated into a single hemisphere at E12.5 and brains were hybridized with HCR probes against *EfnA5* mRNA at P7. Sections were also probed for *Rorβ* mRNA to mark the barrel cortex and segregate superficial (L2/3, L4) and deep (L5, L6) layers. A marked reduction of *EfnA5* is apparent in electroporated S1 (D) compared to the non-electroporated side (E). High-magnification images of deep layer neurons showing GFP+ (and hence *shRNA* expressing) neurons with low or no *EfnA5* signal (F, yellow arrowheads in F') compared with GFP- neurons in the same cortex (F, magenta arrowheads in F'). No GFP+ neurons are visible in deep layers of the non-electroporated side (G), and all neurons show high levels of *EfnA5.*

ns, not significant; *, p<0.05.

Scale bars: 2000 µm (B), 500 µm (B'), 100 µm (E), 25 µm (G').
